# A molecular representation system with a common reference frame for natural products pathway discovery and structural diversity tasks

**DOI:** 10.1101/2024.10.01.616173

**Authors:** Nicole Babineau, Le Thanh Dien Nguyen, Davis Mathieu, Clint McCue, Nicholas Schlecht, Taylor Abrahamson, Björn Hamberger, Lucas Busta

## Abstract

Researchers have uncovered hundreds of thousands of natural products, many of which contribute to medicine, materials, and agriculture. However, missing knowledge of the biosynthetic pathways to these products hinders their expanded use. Nucleotide sequencing is key in pathway elucidation efforts, and analyses of natural products’ molecular structures, though seldom discussed explicitly, also play an important role by suggesting hypothetical pathways for testing. Structural analyses are also important in drug discovery, where many molecular representation systems – methods of representing molecular structures in a computer-friendly format – have been developed. Unfortunately, pathway elucidation investigations seldom use these representation systems. This gap is likely because those systems are primarily built to document molecular connectivity and topology, rather than the absolute positions of bonds and atoms in a common reference frame, the latter of which enables chemical structures to be connected with potential underlying biosynthetic steps. Here, we present a unique molecular representation system built around a common reference frame. We tested this system using triterpenoid structures as a case study and explored the system’s applications in biosynthesis and structural diversity tasks. The common reference frame system can identify structural regions of high or low variability on the scale of atoms and bonds and enable hierarchical clustering that is closely connected to underlying biosynthesis. Combined with phylogenetic distribution information, the system illuminates distinct sources of structural variability, such as different enzyme families operating in the same pathway. These characteristics outline the potential of common reference frame molecular representation systems to support large-scale pathway elucidation efforts.

**Significance Statement:** Studying natural products and their biosynthetic pathways aids in identifying, characterizing, and developing new therapeutics, materials, and biotechnologies. Analyzing chemical structures is key to understanding biosynthesis and such analyses enhance pathway elucidation efforts, but few molecular representation systems have been designed with biosynthesis in mind. This study developed a new molecular representation system using a common reference frame, identifying corresponding atoms and bonds across many chemical structures. This system revealed hotspots and dimensions of variation in chemical structures, distinct overall structural groups, and parallels between molecules’ structural features and underlying biosynthesis. More widespread use of common reference frame molecular representation systems could hasten pathway elucidation efforts.

## 1. Introduction

Natural products are an important source of inspiration for therapeutic compounds (1–3) and are vital in the context of drug discovery (4–7). Due to their structural complexity and frequently low natural abundance, there is great interest and potential in reconstructing natural product biosynthetic pathways in engineered, scalable bioproduction systems (8–10). Such engineering requires identifying biosynthetic pathway genes encoding enzyme catalysts that convert ubiquitous precursors into the complex final products. Almost without exception, these pathway elucidation efforts begin with comparisons of the target natural product’s chemical structure with the structures of co-occurring compounds. Such comparisons greatly assist in developing hypothetical biosynthetic pathways for subsequent testing. Interestingly, the structural comparison step of such projects receives relatively little attention when compared with the emphasis placed on the analysis of nucleotide sequencing data, such as co-expression analysis. This lack of explicit attention may be because even though there exist a great number of molecular representation systems with which to systematically document (and subsequently compare) the structures of complex molecules (11), such methods are largely optimized for drug discovery-related tasks and not pathway elucidation efforts. Generally, molecular representation systems encode a molecule as a string of characters showing atomic connectivity or encode aspects of a given molecule with a series of numbers describing molecular substructures or atomic xyz coordinates. These methods have enabled, for example, structure similarity searches for the rapid identification of potential drug candidates (12), drug-target interaction prediction (13), and toxicity and binding affinity prediction (14, 15). However, these tools can struggle to generate representations that perform optimally in situations outside drug discovery because they are not designed to support the identification and comparison of corresponding atoms and bonds across a set of related molecules. Such comparisons are important operations in tasks related to biosynthesis because the comparisons can provide insights into the origins of specific structural features or the order of operations on a given biosynthetic pathway. Without molecular representations tailored for such tasks, we risk missing critical insights into the biosynthetic and evolutionary aspects of complex molecules, which hampers our ability to understand and leverage natural products to solve real world problems.

Central to a molecular representation system is the documentation of the atoms in a molecule and the connections between them. An intuitive example is the Simplified Molecular Input Line Entry System (SMILES), in which atoms are listed in an order that corresponds to their connectivity. Another widely used set of systems are fingerprint systems, including MACCS (Molecular ACCess System) keys and Extended-Connectivity Fingerprints, which document the presence/absence or count of specific substructures in a given molecule using fixed-length numerical vectors (16, 17). Other approaches focus on 3D arrangements of molecules in space, rather than substructures, and document the xyz coordinates of a molecule’s atoms and information about the bonds between them, such as .PDB files and .XYZ files. Graph-based representations are also popular, in which a molecule’s atoms are coded as nodes and the bonds connecting them as edges (11). Graph representations can be used in conjunction with Graph Neural Networks, which can capture intricate substructures, such as functional groups and motifs, in abstract numerical vector representations called embeddings (18). Overall, SMILES are intuitive and human-readable, fingerprinting methods are efficient and enable large-scale comparisons, 3D methods capture precise molecular geometries needed for understanding molecular conformations as well as interactions, and graphs detail molecular connectivity and topology enabling property prediction (19, 20), reaction prediction (21, 22), and even molecular generation (23–25). Despite these remarkable capabilities, the focus of these representations is generally on the connectivity and relationships between atoms and bonds in a single molecule rather than on the absolute positions of a molecule’s constituents in a standardized reference frame. The lack of a standardized reference frame is a critical point, because it is precisely that frame which facilitates the inference of biosynthetic steps during pathway elucidation.

Due to the lack of a molecular representation system that places molecules in a common reference frame and the potential for such to be a considerable aid with pathway elucidation efforts, the objective of this work was to explore a reference frame-based representation system and its potential to assist in understanding biosynthesis. Interestingly, structural comparisons can also advance our understanding of the evolution of biosynthetic pathways (26), especially when coupled with a phylogenetic data (27). Accordingly, we included data on how the compounds studied here are distributed across a flowering plant phylogeny, thus expanding the scope of our system’s impact. In summary, we provide a unique framework for documenting and comparing molecular features. This approach not only illustrates how we can fill a gap in molecular documentation that will impact natural products research, but also offers new insights into the structural complexity and evolutionary processes underlying chemical diversity.

## Results and Discussion

This study aimed to document chemical structures in a way that would facilitate their analysis via atom- and bond-level comparisons. We chose triterpenoids reported from plant surface waxes as a case study set of molecules because of (i) their structural complexity and diversity, (ii) their well-documented lineage-specific distribution across the plant kingdom, (iii) their relatively well-understood biosynthesis (28) to serve as a positive control, and (iv) applications of triterpenoids and their derivatives (though not necessarily those found in waxes) as therapeutics (29), vaccine adjuvants (30), and commercial drugs (31). After creating a database of 112 triterpenoid structures, we designed a two-dimensional grid-like template onto which backbone atoms were mapped in an x-y coordinate plane and assigned functional groups (Fig. 1A and B). Then, bonds between atom positions were assigned an orientation (in-plane, out-of-page, or into-page orientation). Using this grid-like system, we first focused on analyses of individual positions within the grid, which identified hotspots of variation in triterpenoid structures (section 2A). We then moved to comparisons of each position in the grid between pairs of molecules, leading to hierarchical clustering directly based on structural characteristics, which assembled triterpenoids into distinct structural groups (section 2B). Next, we examined structural variability across all molecules and grid positions simultaneously using direct, structure-based Multiple Correspondence Analysis (similar to Principal Component Analysis) and ordination, revealing key dimensions of structural variation among the triterpenoids (section 2C). Finally, by integrating our triterpenoid grid template with a plant phylogeny, we examined patterns in the phylogenetic distribution of triterpenoids across plant lineages as well as patterns in the co-occurrence of triterpenoid structures and substructures (section 2D). Throughout, we discuss our grid-like molecular documentation system relative to existing methods and discuss advantages and limitations of our approach.

**Fig. 1.**
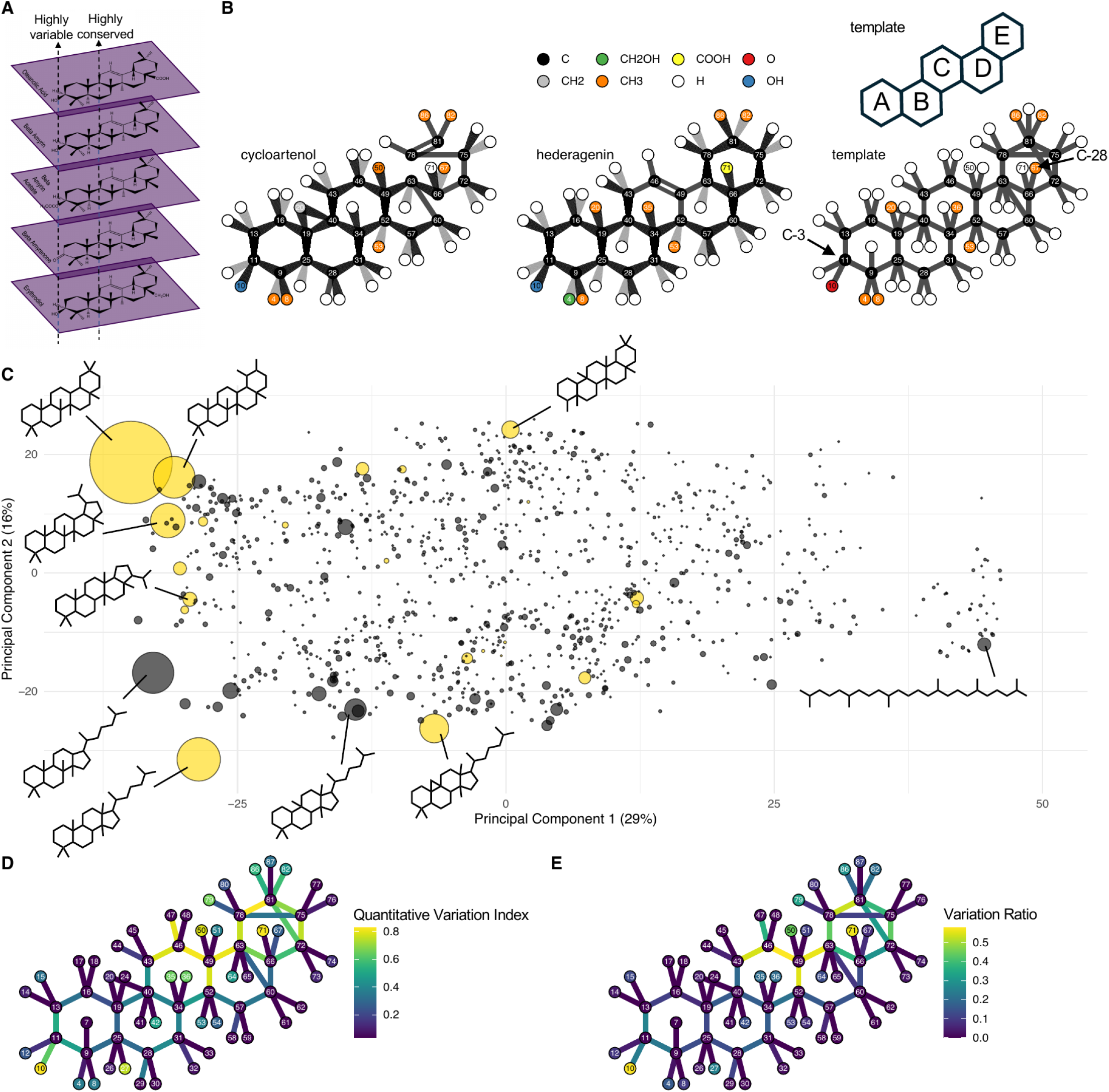
A molecular representation system based on a grid-like template enables the identification of atom-to-atom correspondence. **A** Schematic representation of the atom-to-atom correspondence system. Triterpenoid molecules were aligned to a grid, and corresponding atoms were identified, enabling, for example, identification of positions in the structures that were highly variable versus highly conserved. **B** Example of two structures, cycloartenol and hederagenin, overlayed on our grid template. The template used for all molecules is shown on the right. **C** Principal Component Analysis comparing triterpenoid backbone structures in the TeroKit dataset. Each circle represents a unique skeleton found among TeroKit triterpenoid entries. Circle size corresponds to the number of triterpenoids in the TeroKit dataset with that skeleton, with the largest circle representing 13,434 compounds and the smallest circles representing instances for which there is just a single compound with that skeleton. Circles in yellow represent scaffolds that are represented by the triterpenoids analyzed in this work. **D** The grid template with each atom position and bond position colored according to its Index of Quantitative Variation. **E** The grid template with each atom position and bond position colored according to its Variation Ratio.

### A. A common reference frame molecular representation system reveals hotspots of variation in chemical structures

Our first goal was to collect a set of representative triterpenoid structures with which to work. We began by systematically searching the literature for triterpenoids (C30 compounds) reported from plant cuticular waxes, mixtures of diverse hydrophobic compounds that are found on plant surfaces. 34 reports were identified that, together, reported 581 occurrences of 112 unique triterpenoid compounds on 76 plant species spanning 22 angiosperm plant families. To determine the degree to which these 112 triterpenoids represented overall triterpenoid diversity, we compared the structures of the 112 against the TeroKit database, a collection of >110,000 terpenoid structures (32). First, we extracted the skeletons (carbon connectivity patterns, disregarding stereochemistry and functional groups) of each TeroKit compound, then we filtered for compounds with between C27 and C33 skeletons (39186 unique TeroKit compounds total, and 1075 total skeletons). Next, we computed an Extended-Connectivity Fingerprint for each skeleton, determined pairwise similarity for each pair of skeletons using the Tanimoto metric, and conducted a Principal Component Analysis to visualize the relationships between the skeletons (Fig. 1C). The first two principal components explained 29% and 16% of the total variance among the TeroKit triterpenoids. These two dimensions appear to be associated with variance in the number of rings present in a molecule and the presence of an uncyclized isoprenyl-derived tail instead of an E-ring, though via this fingerprinting approach it is not possible to determine the exact structural features driving variance in these two dimensions. Next, we compared our 112 triterpenoids against the TeroKit compounds. 86 of our 112 triterpenoids were in TeroKit, and these 86, though they only represented 22 of the 1075 TeroKit skeletons, did represent 5 of the top 6 most common skeletons, and the skeletons covered by our compounds encompassed 67% of all the triterpenoids in TeroKit. Having identified that the 112 triterpenoids reported so far from plant waxes are a reasonable representation of the most common types of triterpenoids, our next objective was to develop a grid system capable of documenting triterpenoid structures with a common reference frame. When examining the triterpenoid structures and their commonalities, we noted that, in our test case of triterpenoids found in cuticular waxes, virtually all the molecules of interest had four or five rings. This allowed us to identify a common orientation for each molecule in three-dimensional space and partially align many of their rings in three-dimensional space; aligning all atoms to a standardized reference frame (Fig. 1A). From this set of partially aligned structures, we developed a generic template onto which each structure could be mapped. This template consisted of A, B, and C rings, a four or five-membered D ring, and an optional four or five-membered E ring. The template allowed molecules with only four rings to be directly compared against those with five (as well as comparisons of molecules with four versus five-membered D and E rings) by aligning the uncyclized carbons of tetracyclic molecules with the corresponding cyclized portions of the other molecules (Fig. 1B). This comparability is a reflection of the common precursor to all of these molecules, 2,3-oxidosqualene. In short, the grid allowed for the identification of atom-to- atom correspondence across the structures of 112 triterpenoids (Supplemental Table 1), setting the stage for subsequent comparative analysis.

Comparative, quantitative analyses often rely on comparisons of continuous variables (as opposed to categorical/discrete variables). However, there are a number of well-established methods for comparing entities that can largely only be described by categorical variables, like the molecular subjects of study here. Two prominent methods for categorical variable comparison are the Index of Qualitative Variation, and the Variation Ratio. The Index of Qualitative Variation is a statistical measure of variability within categorical data, ranging from 0 to 1, that measures the evenness of the distribution of cases across different categories. The value of the Index of Qualitative Variation is 0 when there is no variation, meaning all cases are concentrated within a single category, and reaches its maximum value of 1 when there is maximum variation and the cases are distributed evenly across all categories. The Variation Ratio complements the Index of Qualitative Variation by focusing on the proportion of cases in the most common category relative to the total number of cases (1 - f/N), where f is the frequency of the most common category and N is the total number of cases. This measure ranges from 0, when there is no variation and all cases fall into a single category, to values approaching but less than 1, when the most common category coexists alongside a large number of less common categories. Based on their complimentary nature, we hypothesized that the Index of Qualitative Variation and the Variation Ratio could be useful in examining bond and atom distributions across the triterpenoids documented using our grid system.

To examine bond and atom distributions across the triterpenoids documented using our grid system with the Index of Qualitative Variation and Variation Ratio, we considered the orientation of each bond and the identity of each atom/substituent in each position of the grid across all 112 triterpenoids. The possible states for each bond (absent, present in-plane with page, present with out-of-page orientation, present with into-page orientation) and each atom (absent, present as hydrogen, present as carbon, etc.) were considered as the possible “cases” for each variable from which to calculate the Index of Qualitative Variation and Variation Ratio for each grid position. We then visualized the two metrics by plotting them on template triterpenoid structures (Fig. 1D and E). This revealed several regions of high variation and low variation across the triterpenoids in our database. One region of particularly high variability was the bonds at the top (in our orientation) of the C and D rings, meaning that in our molecule set there is considerable stereochemical and single/double bond variation in that region (Fig. 1D and E). Similarly, bonds in the modular D and E rings exhibited considerable variation, consistent with the presence of triterpenoids with 6-6-6-5, 6-6-6-6-5 and 6-6-6-6-6 ring structures in our dataset. Regarding substituents, several regions had particularly high variability, including carbons C-3 and C-28 (atom positions 10 and 67 in our representation system), which are common sites of P450 oxidation. Thus, the present representation system enables analyses based on atom-to-atom correspondence, providing the ability to recognize regions of high and low variability within a set of molecular structures.

### B. Atom-to-atom comparisons in a common reference frame connect hierarchical clustering with biosynthesis

In addition to enabling the determination of dataset-wide regions of high structural variability at the scale of bonds and atoms/substituents, our common reference frame molecular representation system also enabled pairwise comparisons of all grid positions across all pairs of structures. Thus, the number of differences between each pair of structures were quantitatively assessed to create a distance matrix, which served as the foundation for hierarchical clustering. The clustering process yielded a dendrogram that categorized the triterpenoids into four overall structural groups, based on structural similarity (Fig. 2). Since a natural product’s structure follows from the biosynthetic processes that create it, we hypothesized that the hierarchical clustering analysis we performed would mirror underlying biosynthetic processes. To assess whether our structural analysis aligned with biosynthetic pathways, cation intermediates from 2,3-oxidosqualene cyclization were aligned with regions of the tree in which their products were found (Fig. 2). Inside an oxidosqualene cyclase, 2,3-oxidosqualene can be cyclized in a number of configurations, including those that lead, respectively, to the protosteryl, hopanyl, and lupanyl cations (Fig. 2). Interestingly, products derived from the protosteryl cation, for example, lanosterol, cholesterol, cycloartenol, sitosterol, were clustered together in one clade of the dendrogram. Hereafter, we refer to this clade as the protosteranes and indicate it with purple. Also clustered in the dendrogram were products derived from the hopanyl and lupanyl cations (hopenone, lupeol, betulinic acid, etc.) and their rearrangement structures the fernanyl and germanicyl/taraxasteryl cations, respectively (fernenone, germanicol, taraxasterol, taraxasterone, etc.). We refer to this group as the hopane/lupanes and denote them with the color green. The lupanyl and germanicyl cations can undergo further rearrangement inside an oxidosqualene cyclase to form the ursanyl and oleanyl cations, which give rise to a great number of compounds in our dataset, most with 6-6-6-6-6 ring structures, including alpha- and beta-amyrin, ursolic and oleanolic acid, and related derivatives, which were also clustered in the dendrogram. The double bonds in these compounds are primarily located in the C, D, and E rings. Below, we refer to this group as the ursane/oleananes and indicate them with the color blue in the following figures. Finally, the oleanyl cation can undergo a series of carbocation migrations to form a series of charged structures, including the taraxeryl, multiflorenyl, glutinyl, and friedelanyl cations (34). These cations in turn give rise to compounds such as friedelin, glutinol, and baurenol, along with their derivatives, which were clustered together in the dendrogram. Going forward, we refer to this group as the “taraxane/fridelanes” for simplicity and denote them with the color red in the figures. Thus, patterns in hierarchical clustering schemes derived using a molecular representation system with a common reference frame largely mirror underlying biosynthetic processes, including, in the case of our triterpenoids, the underlying relationships between unreleased enzyme intermediates (triterpenoid carbocations). The parallels between hierarchical clustering and underlying biosynthesis obtained when using a common reference frame system will be stronger than when using other molecular representations systems, since those systems are not designed to enable analyses that leverage atom-to-atom correspondence among the whole set of molecules under study. This advantage, together with the results presented above, suggests that the common reference frame can enable researchers to form well-supported hypotheses about the topology of cascading enzyme reactions (like terpenoid cyclization) without isolating or trapping unstable intermediates. Since natural products’ structures are defined by underlying biosynthesis regimes, this ability to form well-supported hypothesis likely extends to modes of biosynthesis other than cascading enzyme reactions, like multi-step pathways.

**Fig. 2.**
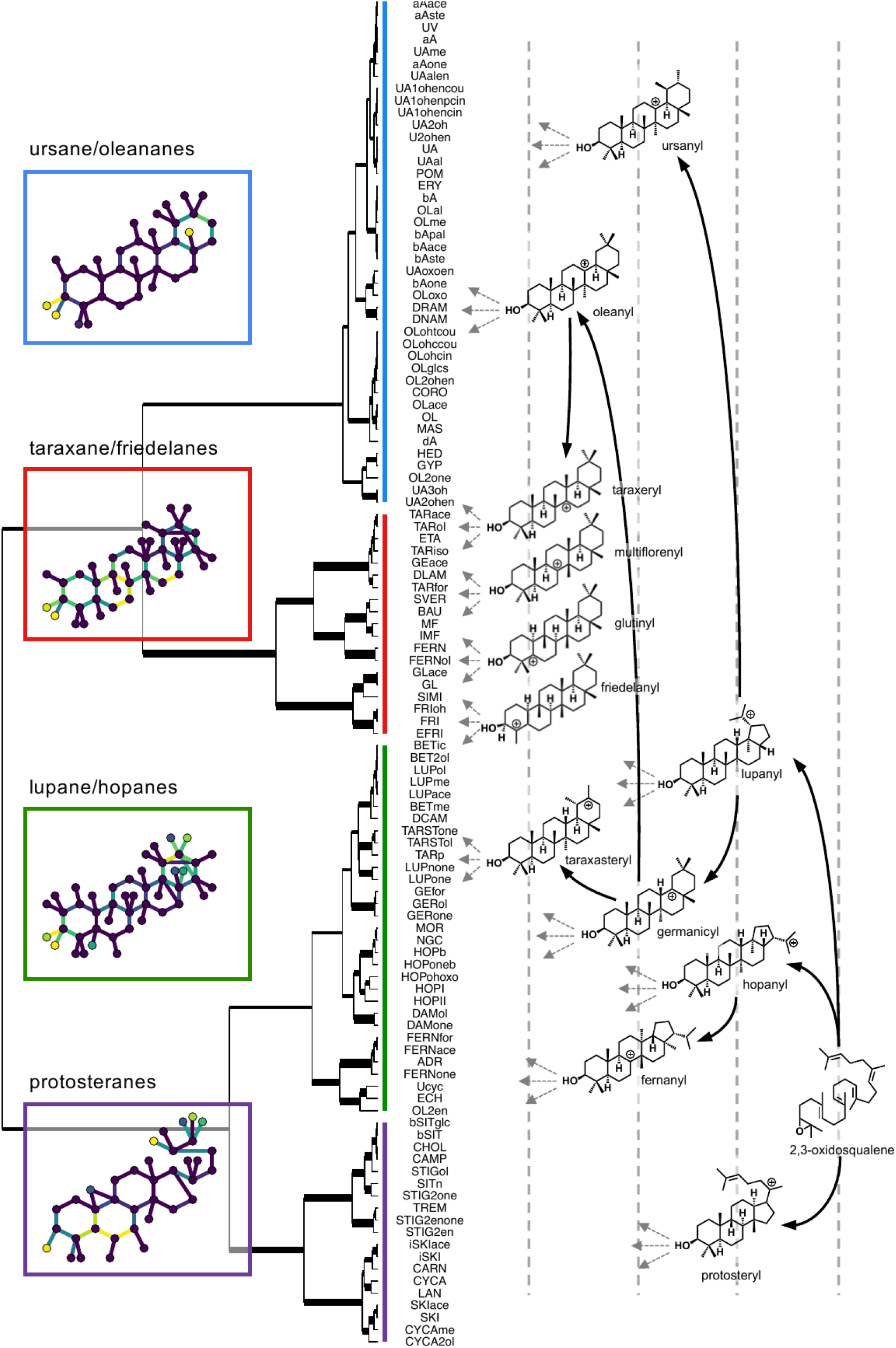
Hierarchical clustering of wax triterpenoid structures and structural variation within each major structural group. Clustering was determined using ward.D clustering on a distance matrix obtained by applying computing Gower distances (33) from a matrix of structural data for the triterpenoids. Four overall structural groups are identified, with the general structures of each shown in inset panels, colors within which represent the Variation Ratio of each bond or atom position within the general structure. Branch widths represent support for respective nodes, with wider widths indicating more support. Cations underlying the biosynthesis of triterpenoids are shown on the right, with the grey dotted lines included simply to guide the eye with regard to the hierarchy of cations. Dotted grey arrows represent further rearrangements to generate the final products shown on the dendrogram. Abbreviations are defined in Supplemental Table 1.

Beyond enabling clustering with an enhanced basis in biosynthesis, a common reference frame representation system allows direct, numerical comparisons of variation at the bond and atom/substituent level across a collection chemical structures. This capability facilitates the application of mathematical analyses typically restricted to numerical datasets. So, having identified four overall groups of triterpenoids in our data set, we next aimed to explore the characteristics of each group through numerical analysis. Within each group, we calculated the Variation Ratio at each bond and atom position in the grid. This revealed that variability in the protosterane group was primarily along the triterpenoid backbone at the bottom of the A and B rings, alongside strong conservation of the stereochemistry of the C and D rings (Fig. 2, purple inset box). Based on the structures of the individual molecules belonging to this group (Supplemental Figure 1), the variation appears to be due to oxidation along the bottom of the A and B rings that changes their stereochemistry. The lupane/hopanes, in contrast, exhibited variation in two major regions, in the bottom left-hand corner of the A-ring (in the present reference frame) and in the top right-hand corner of the E-ring, alongside relatively strong conservation in the arrangement of the B, C, and D rings. These two regions of variation seem likely due to, respectively, modifications of the A-ring functional group by transferase as well as redox enzymes (generating the methylated, acetylated, and oxidized derivatives in this group, Supplemental Figure 1), and ring-opening reactions during early stages of oxidosqualene cyclization (conversion of the lupanyl to the germanicyl/taraxasteryl cations). The minimal variation in the B, C, and D rings seems likely due to the relatively small distance the carbocation migrates across the backbone structure during the generation of these compounds’ precursor cations. The ursane/oleananes exhibited remarkably strong conservation in their core ring structure (6-6-6-6-6), with only minor variation in the E-ring stereochemistry, likely due to the difference in methyl group position on the E-rings of the ursanyl and oleanyl cations themselves. However, this group did exhibit remarkable variability with respect to the identity of the substituents on C-3 and C-28 (positions 10 and 67 in our system). Based on the specific structures of the compounds in the ursane/oleananes (Supplemental Figure 1), it seems that a wide variety of redox and transferase reactions generate structural diversity at C-3 in this group. Last, we examined structural variability in the taraxane/friedelinane overall structural group. These compounds arise from substantial cation migration (compare cation position in the ursanyl cation versus the friedelanyl cation), which causes substantial shifts in the stereochemistry of the backbone carbons causing the variability heat map to light up the A and B rings, as well as the lower portions of the C and D rings (Fig. 2, red inset box). There was also substantial variation in the substituents and stereochemistry around the C-3 carbon (lower left-hand portion of the A-ring), arising from the differences in that region between the friedelanyl-derived compounds versus the other compounds in that group. Thus, the regions of variability among the four overall structural groups appear to stem from at least two sources: variability in oxidosqualene cyclization and modifications post-cyclization (oxidation, acylation, etc.). Signatures from both these sources of variability can be seen in the variability heat map of the overall structural groups (Fig. 2, inset boxes). This highlights the ability of a common reference frame representation system to discriminate between sources of variability in the molecules beings studied and provides insights into the molecules’ biosynthesis based on their structures.

### C. Common reference frame representations enable illustration of major dimensions of structural diversity across many molecules

Having identified that hierarchical clustering using the atom-to-atom correspondence system was capable of identifying regions of variability within overall structural groups in the dataset, our next goal was to explore the potential of analyses that considered variation in the entire dataset as a whole. Often, major sources of variability in a large data set are explored using dimensionality reduction techniques such as Principal Component Analysis that generate eigenvalues for each parameter that correspond to the amount of variability in the data set contained within that parameter. In our case, however, the parameters under consideration were not numerical, such as those used in Principal Component Analysis, but rather categorical. Accordingly, we chose to analyze dimensions of variability in our data set using Multiple Correspondence Analysis, which is analogous to Principal Component Analysis except that it can accept categorical parameters as inputs. To conduct the Multiple Correspondence Analysis, we created a matrix in which each column was a parameter (an atom position or a bond position within the grid) and each row was a molecule. This matrix was passed to the Multiple Correspondence Analysis function which returned three major types of information. The first type of information returned by the Multiple Correspondence Analysis was the amount of variance in the data set explained by each of the dimensions resulting from the analysis (Fig. 3A). Interestingly, the first two dimensions each explained more than 12% of the variance in the data set while the third principal component explained less than 11%. This indicated that there were two major dimensions of variability in our triterpenoid data set followed by a larger number of minor dimensions of variability.

**Fig. 3.**
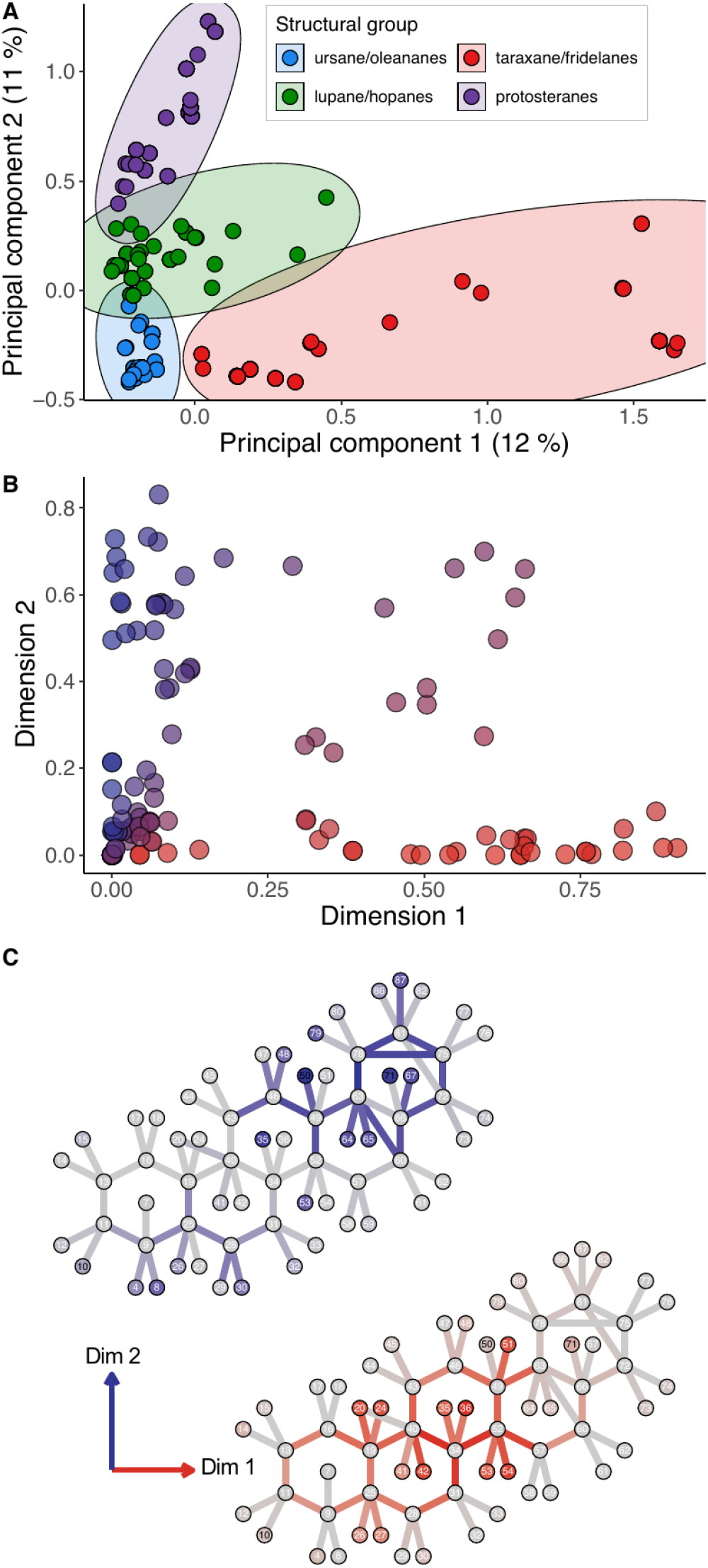
A common reference frame molecular representation system reveals major dimensions of structural variability in triterpenoids from plant surface waxes. **A** Multiple Correspondence Analysis of triterpenoid structures. **B** Ordination output from the Multiple Correspondence Analysis. Each circle represents a bond or atom position in the grid system. **C** Ordination output mapped onto the template grid system. Grey represents bond or atom positions with low contribution to variation in respective dimensions, while a darker color (red for dimension 1, blue for dimension 2) indicates a bond or atom position that contribute substantially to variation in that dimension.

To explore the two major dimensions of variability identified by the Multiple Correspondence Analysis in more detail, we next examined the second type of information provided by the analysis: the position of each molecule in our data set in a space defined by those two major dimensions. That first dimension, plotted along the x-axis, separated one overall structural group, the taraxane/friedelinanes from the other structures (Fig. 3A, discussed in detail below). The second major dimension of variability, plotted along the y-axis, separated the other three overall structural groups, the protosteranes, ursane/oleananes, and lupane/hopanes. In addition to providing information about the position of the different overall structural groups within the two major dimensions of variability in our triterpenoid data set, the Multiple Correspondence Analysis also provided a third type of information: that about portions of the molecules that contributed to variation in each of the two major dimensions. To examine how structural variation in specific molecular regions contributed to major overall dimensions of variability in the data set, we visualized this third type of information, the ordination output, from the Multiple Correspondence Analysis. For this, we plotted each atom and bond position in a two-dimensional space according to its level of contribution to the two major dimensions of variability identified by the Multiple Correspondence Analysis (Fig. 3C). Thus, points in the ordination plot (Fig. 3B) with large X coordinates and small Y coordinates correspond to atom or bond positions whose variance is correlated and associated with a molecule’s variability along principal component one (Fig. 3C). In contrast, points with large Y coordinates and small X coordinates correspond to atom or bond positions whose variance is correlated and varies along principal component two. Points that have both large X and Y coordinates correspond to atom or bond positions that contribute to variation along both the first two dimensions.

Overall, the Multiple Correspondence Analysis, including the ordination component, revealed several interesting trends. First, it showed that the largest source of variability in the dataset (where largest means the greatest number of changes in the identity of the atom at each position in the template or changes in bond orientation) were from variability in the taraxane/friedelinane structural group. Looking back to the hierarchical clustering dendrogram (Fig. 2), this is consistent with the major changes in stereochemistry and methyl group position that accompany the substantial carbocation migration that occurs during the cyclization process leading to these compounds. During these cyclizations, the cation migrates a substantial distance along the compound’s backbone (34), often ending in the A or B ring. This can be seen by the high level of similarity between the heat map of variation for the first dimension (Fig. 3C) with the heat map for variation in the taraxane/fridelane structural group in the dendrogram (Fig. 2, red inset plot). Thus, even though the compounds that make up that structural group are in one relatively small cluster within the dendrogram, they actually represent a substantial portion of the structural diversity within the overall set of molecules, at least, according to the metrics used here. This analysis also emphasizes that molecules derived from the taraxeryl, multiflorenyl, glutinyl, and especially the friedelanyl cation are quite structurally different from other triterpenoids, mainly in their backbone stereochemical structure, which can be easy to overlook when considering these molecules as 2D drawings on a piece of paper or a computer screen.

### D. Atom-to-atom comparisons in a phylogenetic context reveal discrete phases of underlying biosynthetic routines.

Towards the goal of advancing our ability to elucidate biosynthetic pathways, other studies have shown the benefits of considering plant chemical occurrence in a phylogenetic context because doing so can reveal patterns that suggest biosynthetic steps and help us understand the evolution of biosynthetic pathways (27). Accordingly, the present database was used to build Fig. 4, which features a flowering plant phylogeny and triterpenoid occurrence across plant species. Triterpenoid diversity within each species (the number of distinct triterpenoids reported), represented in the right marginal plot, shows that the highest and lowest diversity clusters align with specific plant groups on the phylogenetic tree. For example, *Quercus* (oaks) or *Vaccinium* (the genus to which blueberries belong), have been reported to be rich in triterpenoid diversity and investigators interested in triterpenoid diversity could home in on such lineages

After mapping the triterpenoids in this study onto a simplified plant evolutionary tree, we became interested in how pairs of triterpenoids cooccurred on the species included in that tree, particularly in relation to the pair’s structural similarity. Intuition may suggest the hypothesis that more structurally similar compounds are more likely to co-occur due to shared underlying biosynthetic processes, and the present dataset offered the opportunity to test this hypothesis. For each pair of triterpenoids in our dataset, we computed two metrics: one metric for structural similarity and one metric for whether they appeared together more or less frequently than when they were distributed randomly across the phylogeny. As a metric for structural similarity, we used the proportion of atoms or bonds not shared by the pair of molecules, computed directly from their representations in the common reference frame molecular representation system we are describing here. To determine whether each pair occurred together more or less frequently than expected, we took the number of occurrences of each of the two compounds in each pair and randomized those occurrences across the phylogeny 100 times. We then computed the mean number of times the two compounds cooccurred and the 95 percent confidence interval for that mean. If the actual number of co-occurrences for a pair fell outside the 95% confidence interval for that pair, we considered the pair to have some mutually exclusive or mutually inclusive character. We plotted mutually exclusive and inclusive pairs based on their chemical similarity (x-axis) and the difference between expected and observed co-occurrence (y-axis; Fig. 5A). Many pairs of triterpenoids that co-occurred belonged to the same overall structural group, for example, alpha and beta-amyrin, ursolic acid and oleanolic acid, or campesterol and stigmasterol. This pattern could be due to (i) gene duplication followed by catalytic, but not regulatory divergence, or (ii) enzymes that produce several slightly different compounds, like alpha and beta-amyrin. However, not all pairs of triterpenoids that co-occurred more frequently than expected were structurally related. For example, beta-amyrin and friedelin were found together much more often than expected, even though their structures are quite different. This could be for two reasons: (i) although the biosynthesis of these two compounds is quite similar (friedelin differs from beta-amyrin mainly due to a longer series of carbocation shifts), these shifts create big differences in the 3D arrangement of atoms, but not in how they are connected; and (ii) friedelin synthases have evolved from beta-amyrin synthases multiple times, so they might appear together and be expressed in the same genomes. Although these two factors are likely involved, we cannot rule out the possibility that the higher-than-expected co-occurrence might be because these compounds appear together in well-sampled lineages in our dataset (Fig. 4).

**Fig. 4.**
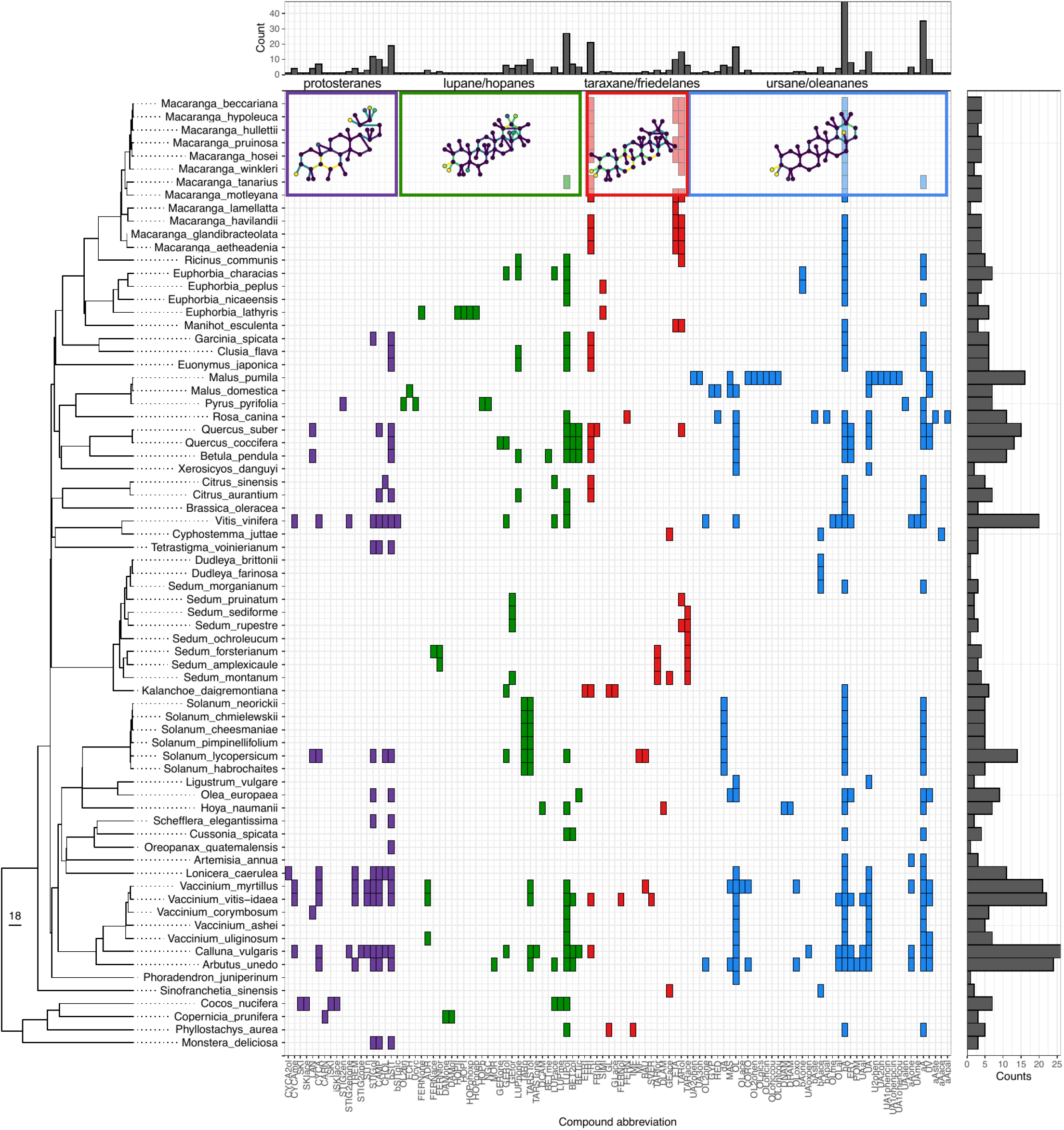
Major triterpenoid structural groups and triterpenoid presence across a plant phylogeny. Our 112 triterpenoid structures from literature reports were mapped onto a pruned phylogeny derived from a previous report (35), based on occurrence within specific plant species. Compound names are listed on the bottom horizontal axis, where each box in the plot represents an instance of that triterpenoid being reported from the cuticular wax mixture of a given plant species (listed on the left vertical axis). Triterpenoid molecules are organized into colored groups based on major structural groups identified by our analysis in Fig. 2; the ursane/oleananes group is purple, the taraxane/friedelanes group is red, the lupane/hopanes group is green, and the protosteranes are purple. The frequency of a triterpenoid compound being reported across all plant species is shown in the top bar chart, while the number of triterpenoid compounds reported in a given plant species is shown in the right bar chart.

**Fig. 5.**
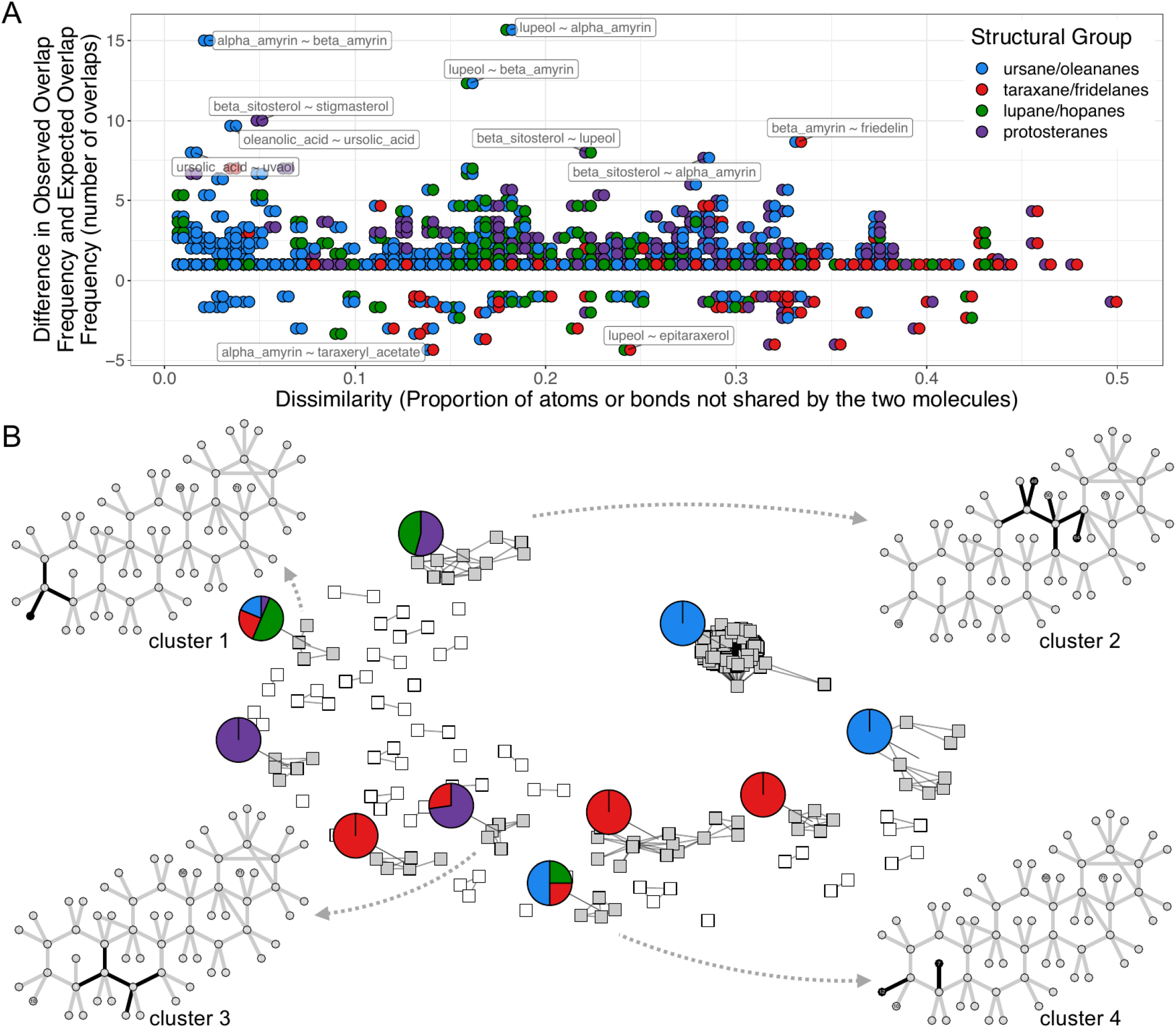
Co-occurrence of triterpenoids as well as atoms and bonds across structurally distinct molecules. **A** Each dot represents a triterpenoid, colored according to structural group as shown in the legend. Triterpenoids are shown as pairs, plotted on the x-axis according to dissimilarity (the proportion of atoms or bonds not shared by the pair), and on the y-axis according to the difference in observed and expected overlap frequency across the phylogeny in Fig. 4. **B** Network in which each node represents a molecular feature (a particular atom in a certain grid position or a bond with a specific orientation in a given grid position). Edges between nodes indicate atoms/bonds that co-occur frequently among molecules in the dataset. Colored clusters grey indicate groups of four or more atoms/bonds that co-occurr with high frequency. Each grey cluster’s pie chart indicate which overall structural groups those co-occurring atoms/bonds are found in.

The above analysis of chemical co-occurrence patterns and chemical similarity can be done with other molecular representation systems, as they also allow for calculating structural similarity. However, the system presented here is unique because it not only calculates similarity but also traces the reason for that similarity or difference directly back to the molecule’s structure. This ability, in turn, lets us study more detailed patterns, like how specific bonds or atom positions co-occur, rather than just looking at whole molecule co-occurrence. To illustrate this capability, we looked at all pairs of bonds or atoms in our dataset of triterpenoids (like CH3 in position 8 appearing alongside CH3 in position 4, or OCH3 in position 10 co-occurring with a double bond between positions 75 and 78, and so on). In total, we analyzed 140250 pairs of atoms or bonds and counted how many times they appeared together in the molecules of our dataset (Fig. 5B). Not surprisingly, most pairs of atoms or bonds never appeared together, but a few did co-occur often in the 112 molecules we studied. To account for the varying occurrence rates of individual atoms and bonds, we normalized the frequency of co-occurrence of each pair of analytes to the sum of their individual occurrences minus the co-occurrence count itself (Fig. 5C). This metric gives a relative measure of how often the two atoms or bonds co-occur in a given molecule relative to the number of overall appearances of those individual atoms or bonds in the dataset. We then set a threshold for that metric (0.9), above which we considered the pair to be co-occurring at a high rate. We visualized the relationships between highly co-occurring pairs of atoms/bonds using a network diagram, then used connectivity within the network to identify clusters of highly co-occurring atoms/bonds (grey boxes, Fig. 5B). Next, we determined which overall structural groups (colored clades in Fig. 2) contained the triterpenoids in which each highly co-occurring cluster of atoms/bonds was found (pie charts, Fig. 5B). Many clusters contained atoms/bond pairs that co-occurred frequently within a single overall structural group. However, there were four clusters of highly co-occurring atoms/bonds that were found in triterpenoids that belonged to *multiple* overall structural groups (clusters from which dotted arrows derive, Fig. 5B) This finding was highly interesting, because it means that a common reference frame system enables the identification of sets of bonds/atoms that co-occur quite often in molecules that are otherwise structurally distinct.

We next followed up in detail on each cluster of atoms/bonds that co-occurred with high frequency across otherwise structurally distinct molecules (i.e. the clusters from which dotted arrows start,5B). Cluster 1 was defined by an oxygen in position 10, in-plane single bonds between positions 11-13 and 9-11, and a double bond between position 10 and 11. This set of atoms/bonds was found in 16 different molecules from across all four overall structural groups identified in Fig. 2 (for example, germanicone, beta-amyrinone, fernenone, and stigmastane-3-6-dione). In simple terms, this wide-spread, highly-cooccurring cluster of atoms/bonds could be called ‘an oxo group on C-3’ (position 10 in our representation system). Previous research has shown that this sort of oxo group can arise from P450 oxidase activity (36), suggesting that the biosynthesis of the other compounds in our dataset that share this set of atoms/bonds also arise, in part, from P450 activity (with the exception of friedelin and likely hydroxy-friedelin, in which the oxo group arises directly from oxidosqualene cyclase activity (37, 38)). Thus, our analysis strongly suggests that there are P450s operating in the lineages producing the molecules with C-3 oxo groups. Furthermore, since many of these compounds, including representatives from distinct overall structural groups, co-occur in the same species (Fig. 4) it seems likely that some of these P450s may be promiscuous with respect to their substrates, or that these species may harbor multiple C-3 oxidases with distinct substrate specificities.

Clusters 2 and 3 consisted of 35 and 11 molecules, respectively. The common atom/bond combination within each group was a particular type of backbone structure: an out-of-plane stereochemistry at the linkage between the C- and D-rings (cluster 2), and a double bond in the B-ring (cluster 3). The atom/bond combinations of each cluster were shared across molecules from multiple overall structural groups, suggesting that there are biosynthetic commonalities to backbone structure formation among otherwise structurally distinct molecules. Finally, we examined cluster 4, which included in-plane single bonds between the carbons in positions 10 and 11 as well as positions 7 and 9, together with a methoxy group in position 10, an oxygen in position 12 and a hydrogen in position 7. This combination of atoms/bonds were found in 4 compounds representing three distinct overall structural groups (methyl 3,4-seco- urs-12-en-3-oate (DRAM), methyl 4(23)-dihydrolacunosic acid (DLAM), methyl 4(23)-dihydronyctanthic acid (DNAM), and methyl dihydrocanaric acid (DCAM), reported from the Apocyanaceous epiphyte *Hoya naumanii* (39)). Interestingly, these compounds belong to two different overall structural groups, which, drawing from the example of the C-3 oxo group discussed above (position 10 in our system), suggests that *H. naumanii* bears one or more P450 oxidases with A-ring regiospecificity that potentially accept more than one substrate.

In summary, our analyses show that a molecular representation system built with a common reference frame allows detailed comparisons of chemical structures at the level of atoms and bonds. These comparisons include assessments of atom and bond co-occurrence that in turn provide novel insight into the biosynthetic processes underlying the represented molecules (the comparison of Cluster 1 and Cluster 4, above). Here we used such comparisons to highlight the presence of multi-step biosynthetic pathways in which one enzyme class generates structurally diverse compounds that are further transformed by a second class of enzyme. Our analyses further suggest that these pathways contain enzymes with relaxed substrate specificity.

## 3. Conclusion

Here, we explored the use of a molecular representation system with a common reference frame for tasks related to natural products structural diversity and biosynthesis. We manually entered 112 triterpenoid molecules into the system and made several major findings that were possible only because the system enabled analyses that relied on the ability to identify corresponding bonds and atoms/substituents across the entire set of molecules. First, it was possible to directly measure variation in the molecules’ backbone stereochemistry and attached chemical groups, revealing positions in their general structure that are highly variable or highly conserved. In addition, the common reference frame representation system supported hierarchical clustering that not only identified overall structural groups that each arise largely from similar starting compounds. The system also allowed us to compare the similarities and differences within these groups, down to the level of atoms, chemical groups, and bonds. Our system also worked with tools that simplify complex data, such as Multiple Correspondence Analysis, which helped us find key patterns in how the structures varied. For the triterpenoids we studied, these patterns were linked to changes that occur during the chemical process of oxidosqualene cyclization. Finally, the level of detail in our system, along with data about how the studied compounds are distributed across species, gave us insights into how they are made. It showed cases of (i) likely gene duplication and neofuntionalization, (ii) different molecules formed by the same multifunctional enzyme, and (iii) pathways where a variety of compounds can be changed in similar ways by a single enzyme or enzyme type. The common reference frame molecular representation system presented here has a few limitations in its current form. First, this type of system likely only works for compounds that come from similar starting materials, so that a common template can be created. Automating the process of adding molecules into the system may take some effort to make sure the shared reference frame is maintained. Here, we manually entered each triterpenoid into our system, which was a time-consuming task. However, our work shows the potential benefits of using a common reference system for studying natural product biosynthesis, justifying and supporting future efforts to develop methods that create these systems, especially methods that can be partially or fully automated using existing chemical databases. For one example of such a method, please see the companion to this paper (Mathieu et al., 2024), in which molecular skeletons are used as a templates for common reference frames, enabling the analysis of tens of thousands of molecules.

## 4. Methods

### A. Diversity metrics and matrix analyses

To evaluate atoms/bonds using qualitative variation and variation ratio, we defined custom functions (see Supplemental Code), then imported the 112 terpenoids in the grid system (see Supplemental Table 1), grouped the data by each grid position and applied the Qualitative Variation Index and Variation Ratio functions to each group. The output data were merged and visualized using a custom drawMolecules function (see Supplemental Code), which colors the template structure based on the calculated Qualitative Variation Index or Variation Ratio values. To calculate the Coefficient of Unlikeability between pairs of molecules, molecular pairs were aligned by their atom and bond positions, and the coefficient was computed as the proportion of differing positions relative to the total positions. The resulting data was transformed into a distance matrix, which was subsequently used for additional analyses. Principal Component Analysis was performed on a distance matrix derived from the Tanimoto similarities of molecular fingerprints, allowing us to visualize the different major triterpenoid structural groups. The Principal Component Analysis was performed using the runMatrixAnalysis wrapper (see Supplemental Code), that leverages the FactoMineR R package function ‘pca’. Similarly, Multiple Correspondence Analysis was performed to explore dimensions of diversity among categorical atoms/bonds. The analysis was executed using the runMatrixAnalysis wrapper, which again leveraged the FactoMineR package function ‘mca’. Hierarchical clustering was conducted on the molecular dataset using the runMatrixAnalysis wrapper, which wraps the ‘hclust’ function from the ‘stats’ R package and uses the ‘daisy’ function from the ‘cluster’ R package to create bootstrapped replicates. Euclidean distances were used, and the ‘ward.D’ was selected as the agglomeration method. Visualization was performed using ggplot. All steps are shown in the complete code provided in the project repository at github.com/thebustalab/common_reference_frame.

### B. Phylogenetic analysis, co-occurrence analysis, and network analysis

A phylogeny for the species present in the dataset was created by pruning an existing plant megaphylogeny (35) using the custom function buildTree (see Supplemental Code). The occurrences of each pair of triterpenoids were then randomly distributed across the phylogeny and the average number of co-occurrences that occurred at random (the null model) was compared against the number of observed occurrences in the actual data set. The differences between observed and expected overlaps were analyzed for significance, and the results were visualized using ggplot2. A molecular feature co-occurrence matrix was constructed that captures the co-occurrence frequencies of analytes across different molecules. To normalize this matrix, each co-occurrence frequency is divided by the sum of the occurrence counts of the two analytes involved, minus the co-occurrence frequency itself. This normalization step adjusts the co-occurrence frequencies relative to the total occurrences of each analyte, ensuring that the resulting values reflect the relative co-occurrence between analytes, independent of their individual occurrence rates. A threshold of 0.9 was used to filter the normalized co-occurrence matrix, which selected for the top 10 percent of most frequently co-ocurring atoms/bonds were selected for further analysis. Using these co-occurrences, a network layout was generated with the Fruchterman-Reingold algorithm, and clustering of network nodes was achieved using DBSCAN, with parameters set to eps = 0.2 and minPts = 3.5. The resulting clusters were analyzed and visualized in conjunction with structural group information, revealing the interconnectedness of atoms/bonds within the dataset.

## 5. Acknowledgements

The authors wish to acknowledge support in the form of startup funds to LB from the Swenson College of Science and Engineering. We also extend our thanks to the University of Minnesota Duluth for supporting NB through the Undergraduate Research Opportunities Program. LETD and CM are grateful for the support of the Department of Chemistry and Biochemistry and the UMD Chemistry Master’s Program. The authors also gratefully acknowledge support from Alan Olyer with manuscript preparation. The project described was supported by Grant Number T32-GM110523 from National Institute of General Medical Sciences of the National Institutes of Health. DM acknowledges funding from an NSF-IMPACTS Training Grant (DGE-1828149), NSF Dimensions of Biodiversity (DEB 1737898), and a Jeff Schell Bayer Foundation Fellowship 2022 (Application Number JS-2022-029).

Finally, we collectively acknowledge that the University of Minnesota Duluth is located on the traditional, ancestral, and contemporary lands of Indigenous people. The University resides on land that was cared for and called home by the Ojibwe people, before them the Dakota and Northern Cheyenne people, and other Native peoples from time immemorial. Ceded by the Ojibwe in an 1854 treaty, this land holds great historical, spiritual, and personal significance for its original stewards, the Native nations, and peoples of this region. We recognize and continually support and advocate for the sovereignty of the Native nations in this territory and beyond. By offering this land acknowledgment, we affirm tribal sovereignty and will work to hold the University of Minnesota Duluth accountable to American Indian peoples and nations.

## 7. Data Availability Statement

All raw data and code used to conduct analysis are provided in the project repository at github.com/thebustalab/common_reference_frame.

## 8. Supplementary Material

Supplemental Table 1: Triterpenoid structures in a common reference frame system. Supplemental Figure 1: The 112 triterpenoid structures used in this study. Supplemental Code: github.com/thebustalab/common_reference_frame.

