## Supplemental Figure 1 for "A molecular representation system with a common reference frame for natural products pathway discovery and structural diversity tasks"

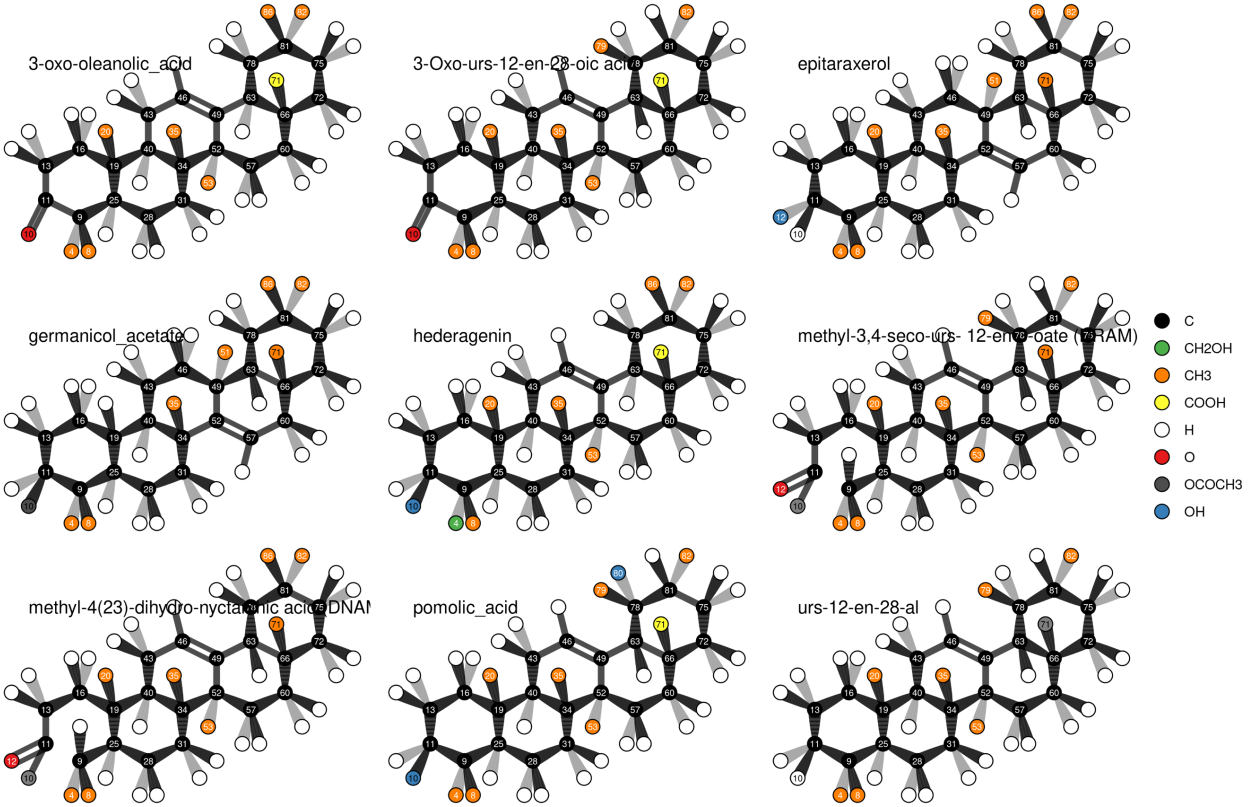
3-oxo-oleanolic_acid
3-Oxo-urs-12-en-28-oic aci
epitaraxerol
germanicol_acetate
hederagenin
methyl-3,4-seco-urs- 12-en -oate ( RAM)
methyl-4(23)-dihydro-nycta nic acid DNA
pomolic_acid
urs-12-en-28-al
C
CH2OH
CH3
COOH
H
O
OCOCH3
OH

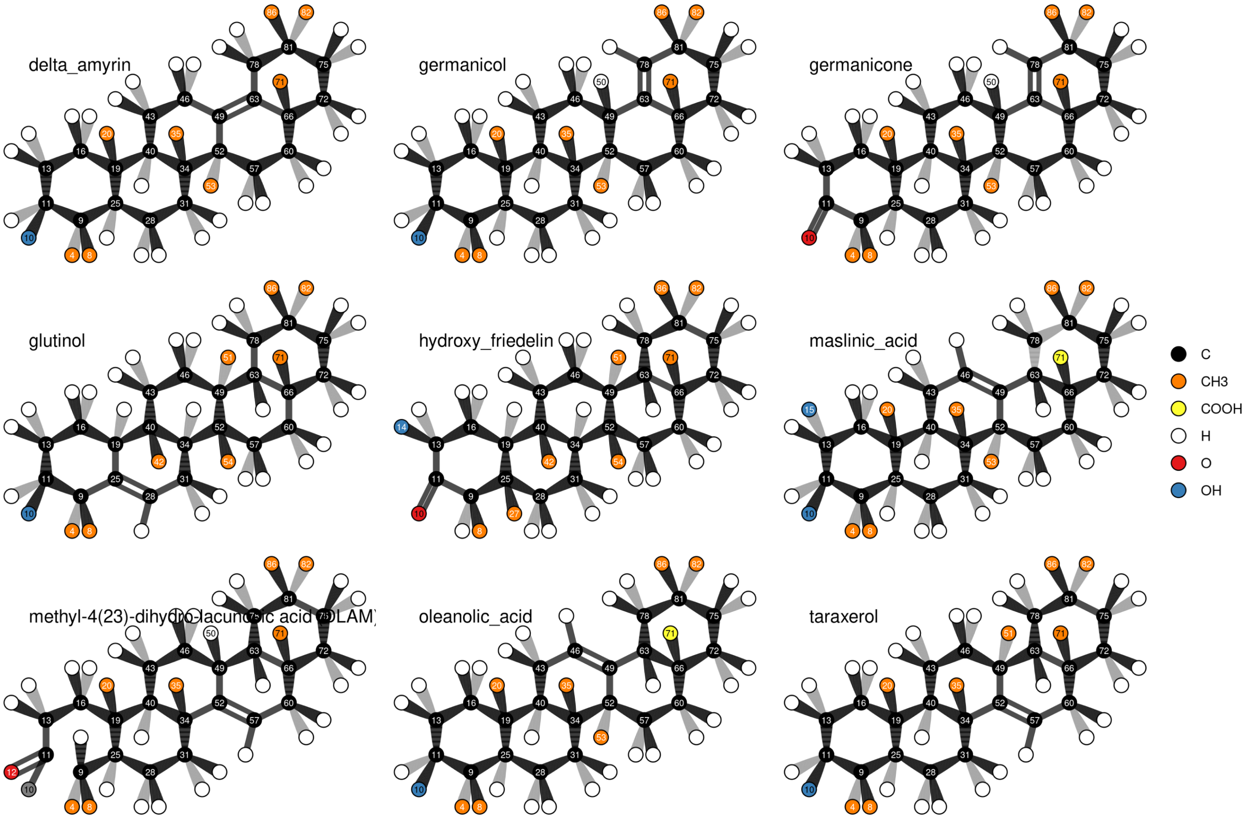
delta_amyrin
germanicol
germanicone
glutinol
hydroxy_friedelin
maslinic_acid
methyl-4(23)-dihydro-lacunosic acid (DLAM)
oleanolic_acid
taraxerol
C
CH3
COOH
H
O
OH

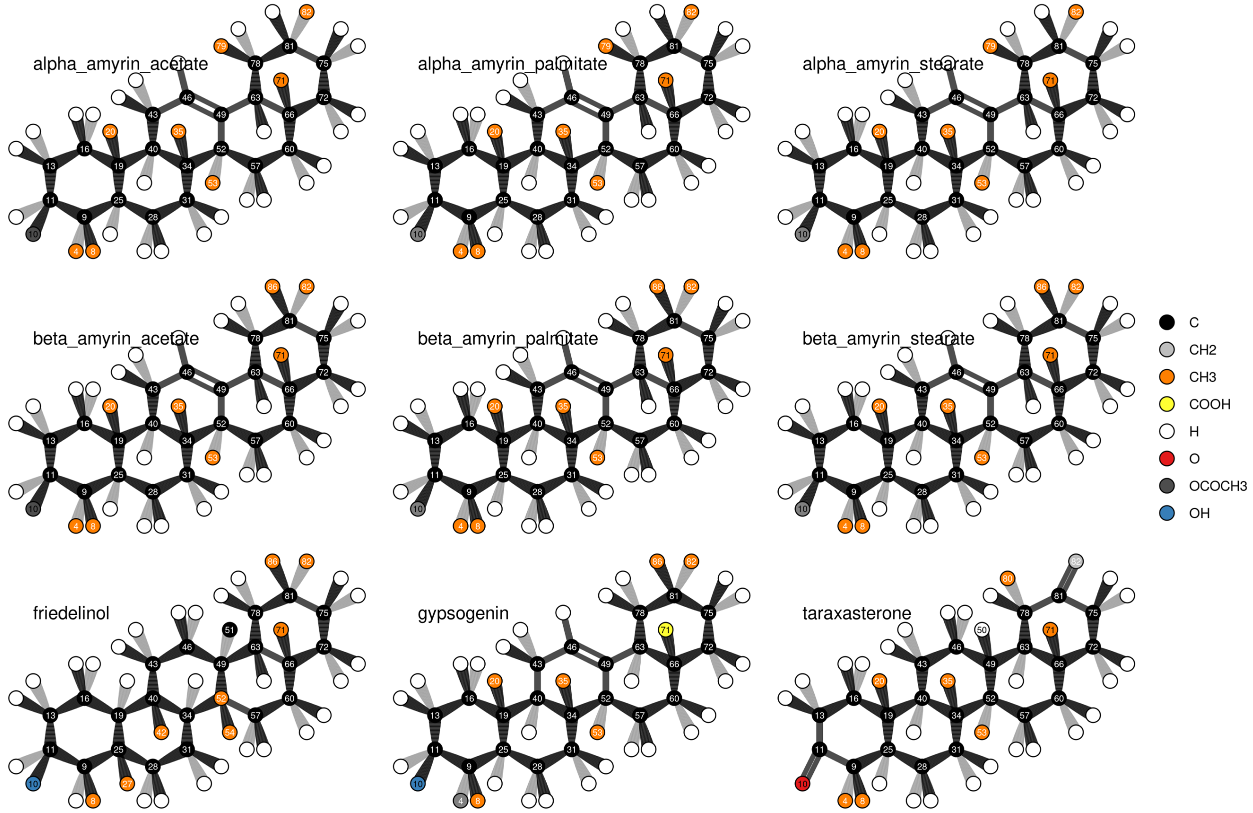
alpha_amyrin_acetate
alpha_amyrin_palmitate
alpha_amyrin_stearate
beta_amyrin_acetate
beta_amyrin_palmitate
beta_amyrin_stearate
friedelinol
gypsogenin
taraxasterone
C
CH2
CH3
COOH
H
O
OCOCH3
OH

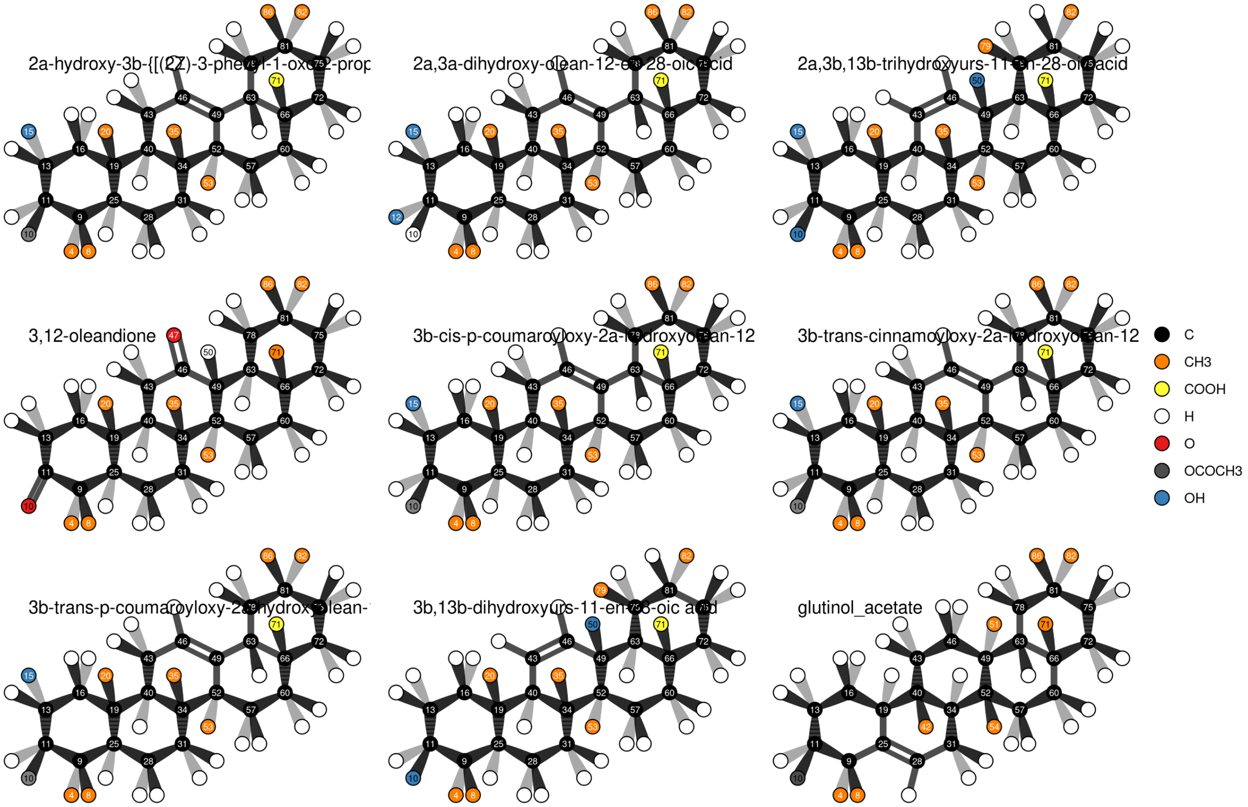

2a-hydroxy-3b-{[(2Z)-3-phenyl-1-oxo-2-prop
2a,3a-dihydroxy-olean-12-en-28-oic acid
2a,3b,13b-trihydroxyurs-11-en-28-oic acid
3,12-oleandione
3b-cis-p-coumaroyloxy-2a-hydroxyolean-12
3b-trans-cinnamoyloxy-2a-hydroxyolean-12
3b-trans-p-coumaroyloxy-2a-hydroxyolean-
3b,13b-dihydroxyurs-11-en-28-oic acid
glutinol_acetate
C
CH3
COOH
H
O
OCOCH3
OH

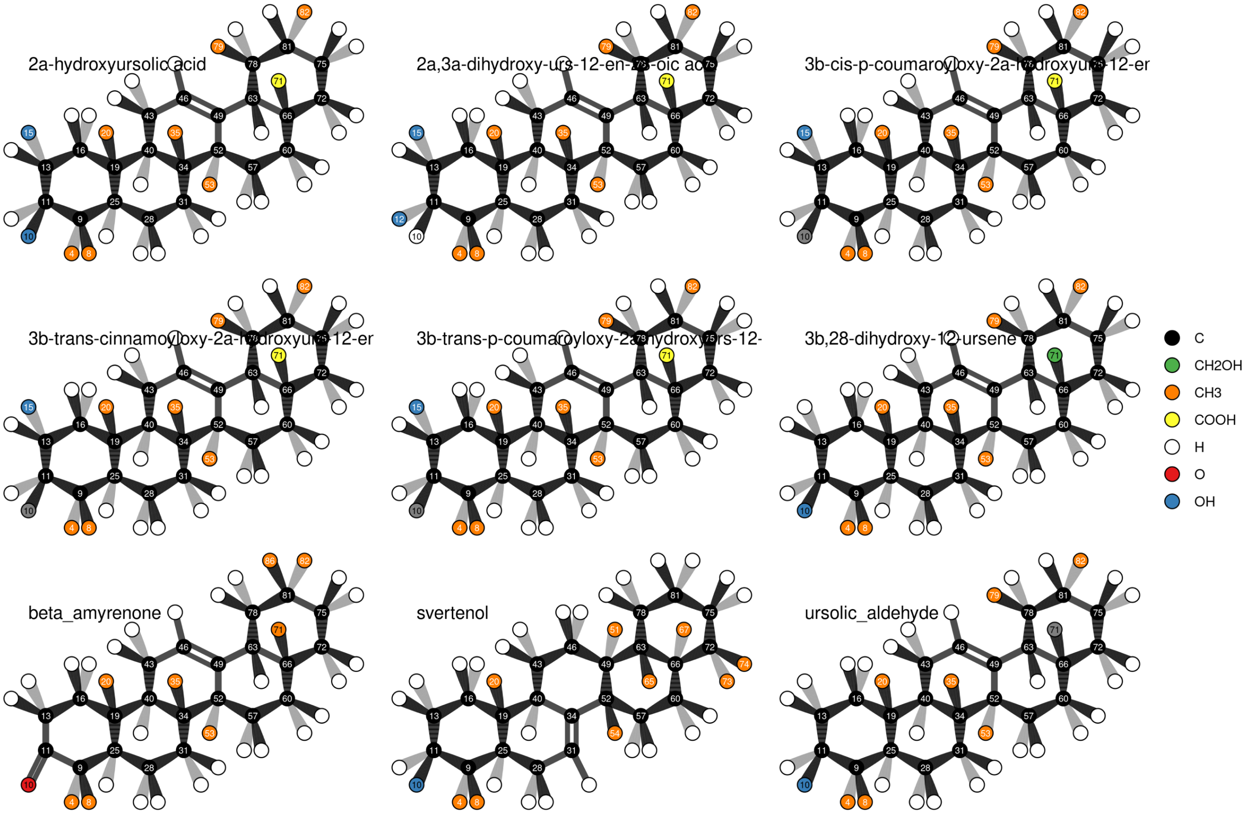
2a-hydroxyursolic acid
3b,28-dihydroxy-12-ursene
beta_amyrenone
svertenol
ursolic_aldehyde
C
CH2OH
CH3
COOH
H
O
OH

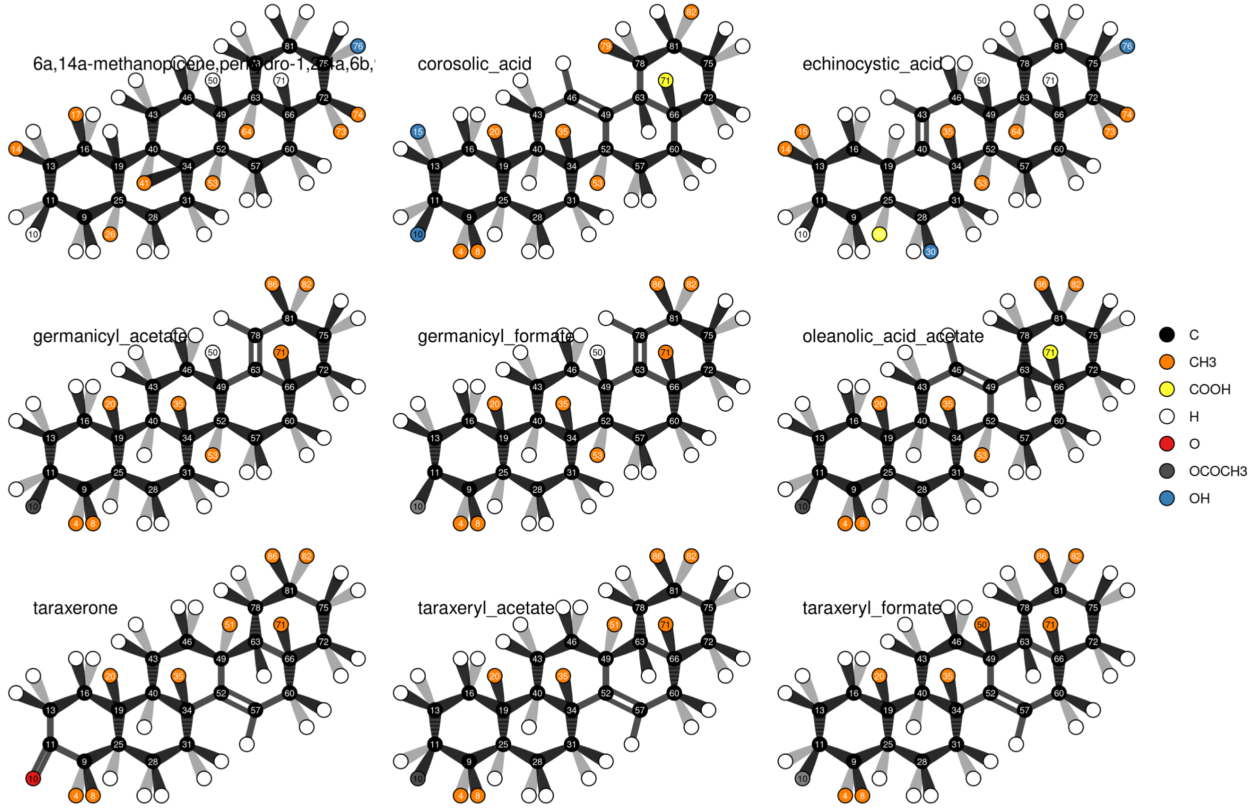
corosolic_acid
echinocystic_acid
germanicyl_acetate
germanicyl_formate
oleanolic_acid_acetate
taraxerone
taraxeryl_acetate
taraxeryl_formate
C
CH3
COOH
H
O
OCOCH3
OH

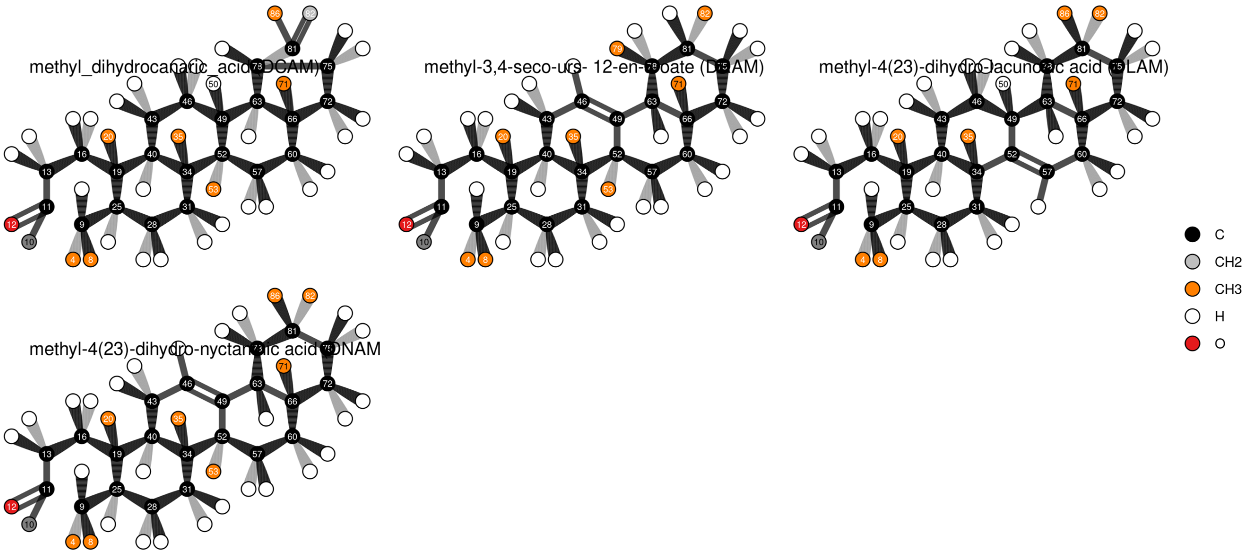

methyl_dihydrocanaric_acid (DCAM)
methyl-3,4-seco-urs- 12-en-oate
methyl-4(23)-dihydro-lacunosic acid
methyl-4(23)-dihydro-nyctanthic acid (DNAM)
C
CH2
CH3
H
O

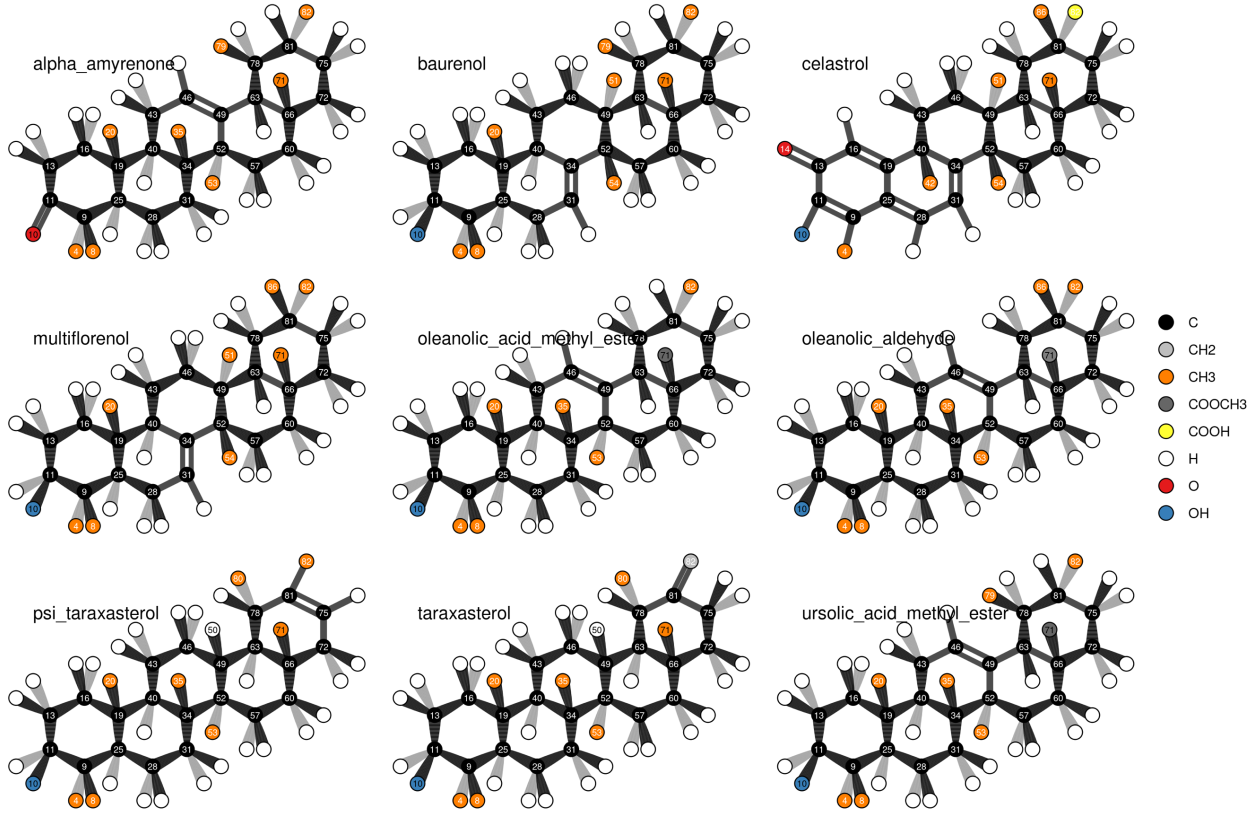
alpha_amyrenone
baurenol
celastrol
multiflorenol
oleanolic_acid_methyl_ester
oleanolic_aldehyde
psi_taraxasterol
taraxasterol
ursolic_acid_methyl_ester
C
CH2
CH3
COOCH3
COOH
H
O
OH

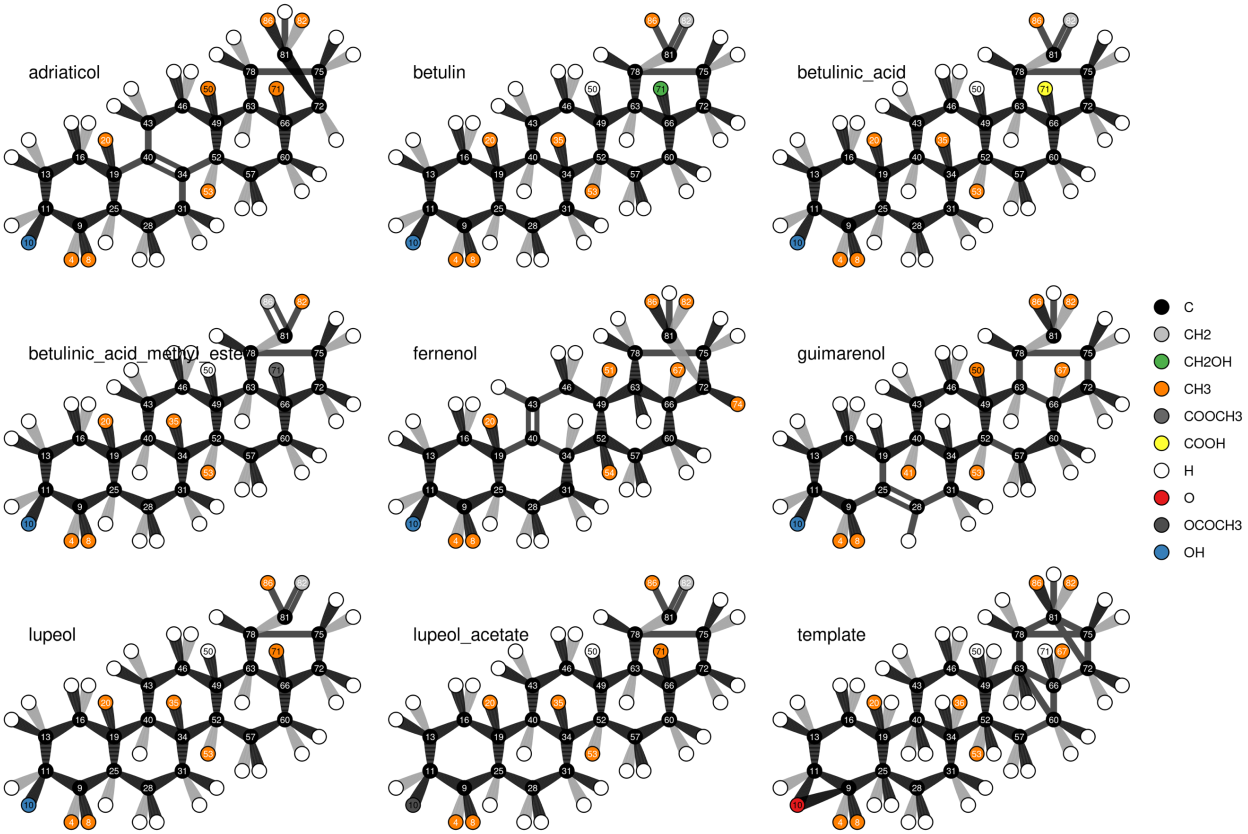
adriaticol
betulin
betulinic_acid
betulinic_acid_methyl_este
fernenol
guimarenol
lupeol
lupeol_acetate
template
C
CH2
CH2OH
CH3
COOCH3
COOH
H
O
OCOCH3
OH

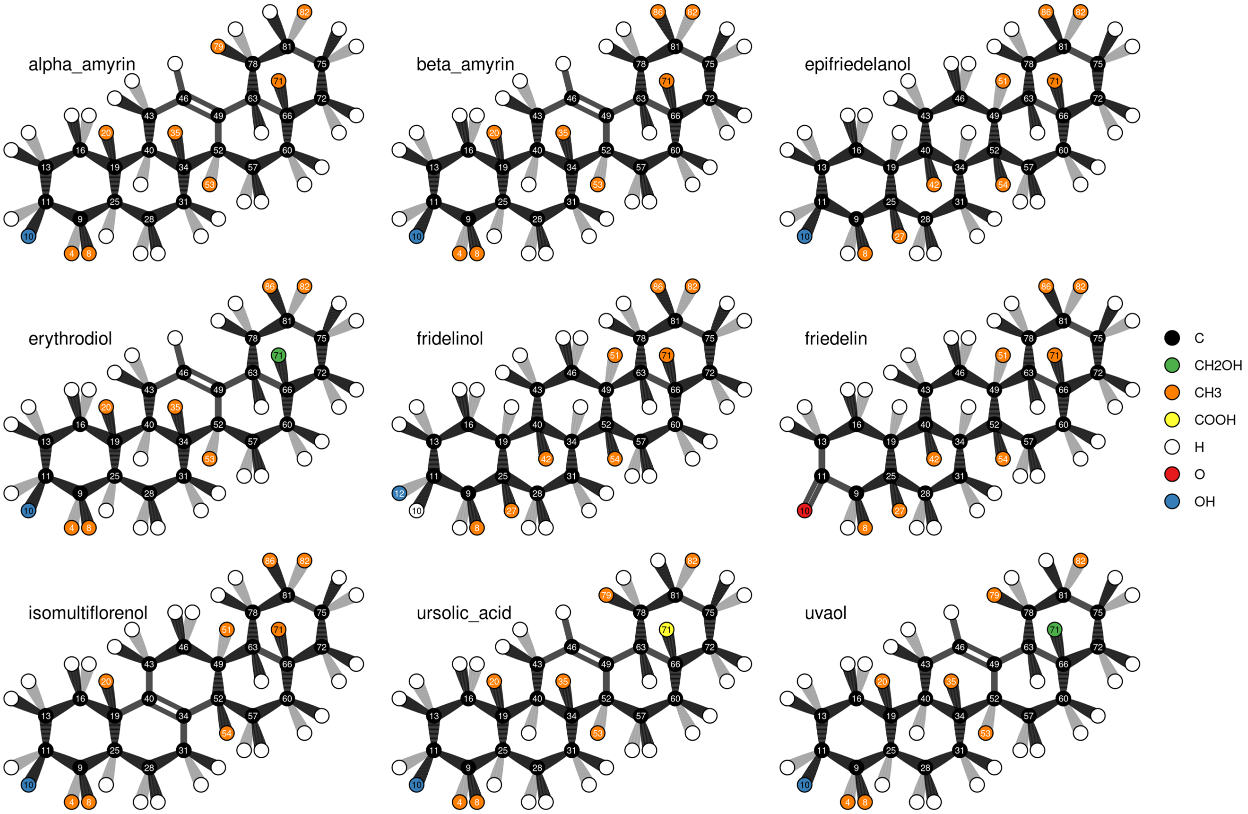
alpha_amyrin
beta_amyrin
epifriedelanol
erythrodiol
fridelinol
friedelin
isomultiflorenol
ursolic_acid
uvaol
C
CH2OH
CH3
COOH
H
O
OH

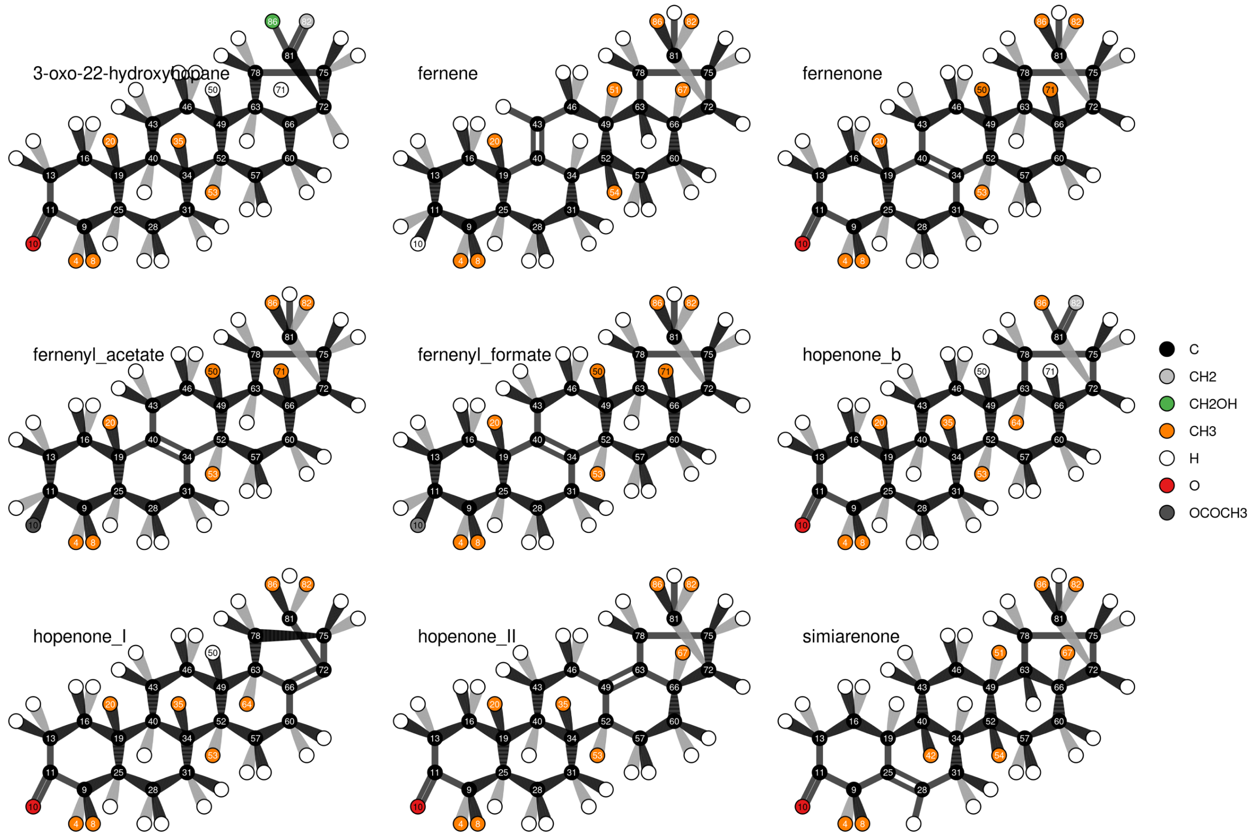
3-oxo-22-hydroxyhopane
fernene
fernenone
fernenyl_acetate
fernenyl_formate
hopenone_b
hopenone_I
hopenone_II
simiarenone
C
CH2
CH2OH
CH3
H
O
OCOCH3

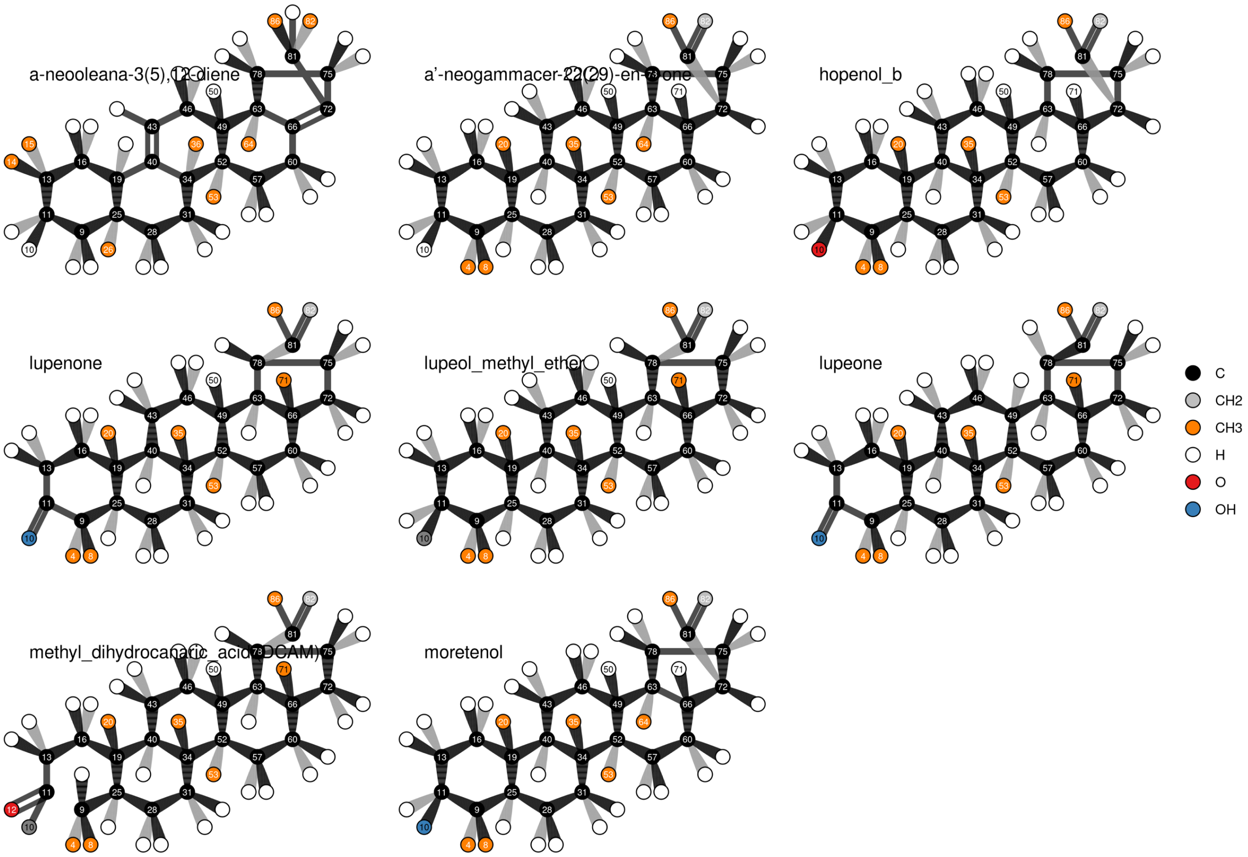
a-neooleana-3(5),12-diene
a'-neogammacer-22(29)-en-3-one
hopenol_b
lupenone
lupeol_methyl_ethe
lupeone
methyl_dihydrocanaric_acid (DCAM)
moretenol
C
CH2
CH3
H
O
OH

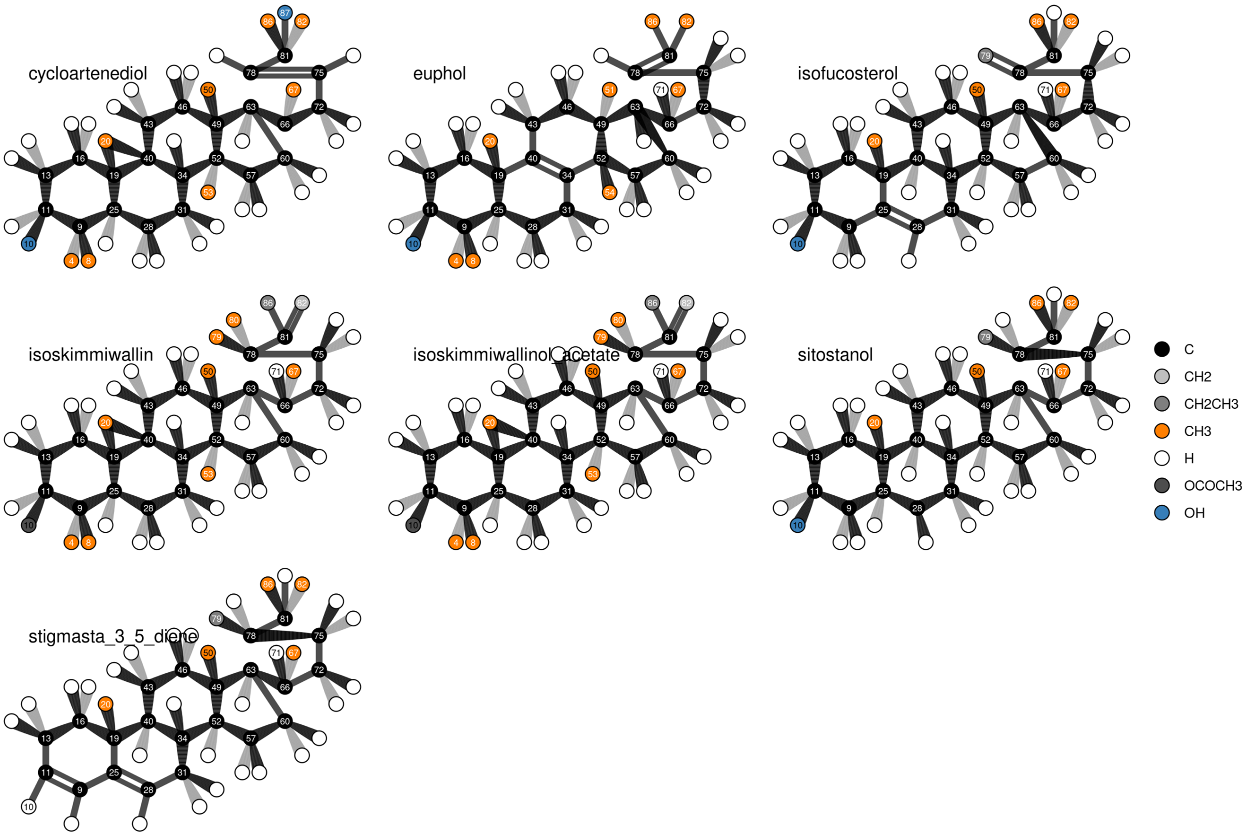
cycloartenediol
euphol
isofucosterol
isoskimmiwallin
isoskimmiwallinol_acetate
sitostanol
stigmasta_3_5_diene
C
CH2
CH2CH3
CH3
H
OCOCH3
OH

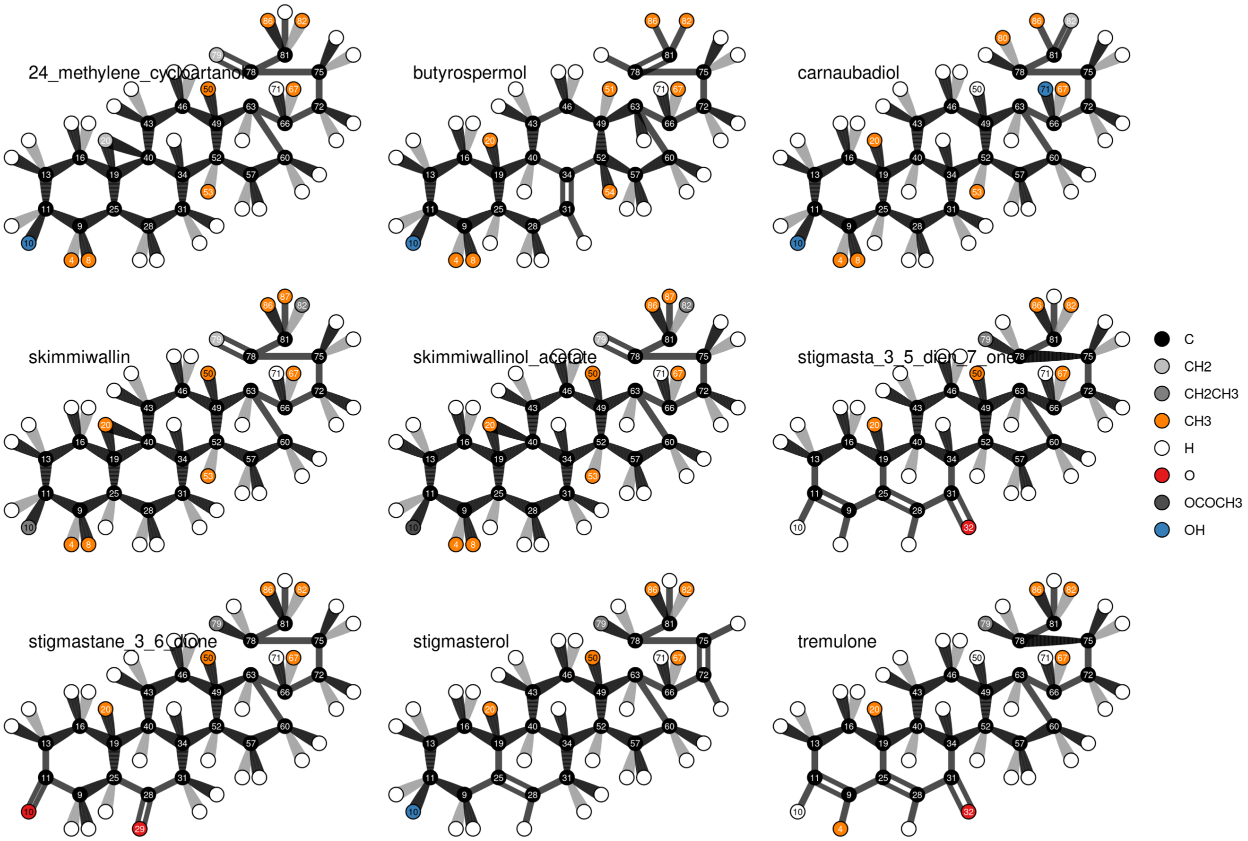
24_methylene_cycloartanol
butyrospermol
carnaubadiol
skimmiwallin
skimmiwallinol_acetate
stigmasta_3_5_dien_7_one
stigmastane_3_6_dione
stigmasterol
tremulone
C
CH2
CH2CH3
CH3
H
O
OCOCH3
OH

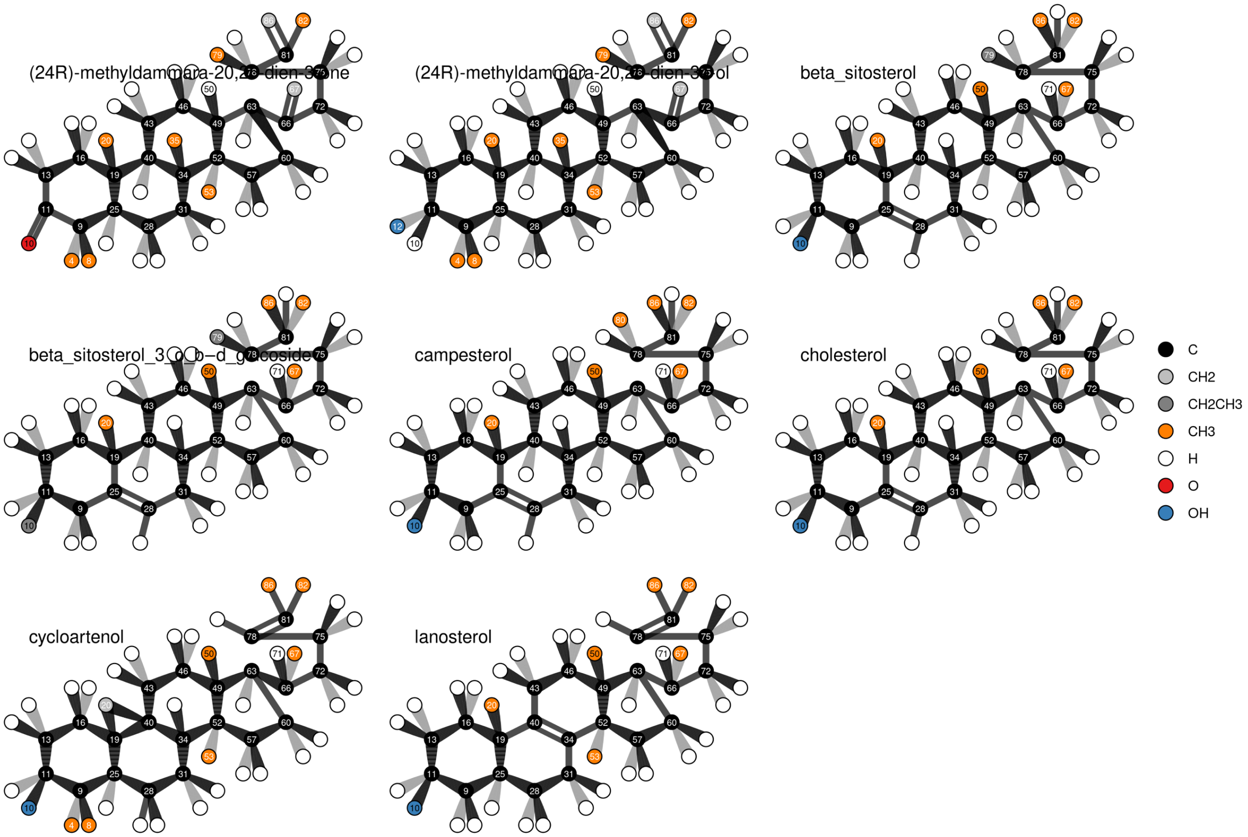
(24R)-methyldammara-20,2[illegible]-dien-3-one
(24R)-methyldammara-20,2[illegible]-dien-3-ol
beta_sitosterol
beta_sitosterol_3_o_b-d_glucoside
campesterol
cholesterol
cycloartenol
lanosterol
C
CH2
CH2CH3
CH3
H
O
OH
